# GPNMB overexpression- a marker of resistance to CDK4/6 inhibitors

**DOI:** 10.64898/2026.03.04.709413

**Authors:** Yuan Gu, Ling Ruan, Yuning Hou, Melissa Gilbert-Ross, Thelma Brown, Kevin Kalinsky, Sunil S. Badve, Yesim Gökmen-Polar

## Abstract

Resistance to cyclin-dependent kinase 4/6 inhibitors remains a major clinical challenge in treating estrogen receptor-positive breast cancer, with no reliable predictive biomarkers currently available for patient selection. To investigate resistance mechanisms, we generated drug-tolerant persisters (DTPs) to abemaciclib and palbociclib in a panel of estrogen receptor-positive breast cancer cell lines. Functional analyses revealed that DTPs showed resistance to CDK4/6 inhibition, maintained G1 arrest, and exhibited increased senescence phenotype. To identify clinically relevant markers of resistance, we compared transcriptomic profiles from DTPs with publicly available gene-expression data from the phase III PEARL trial. Glycoprotein non-metastatic B (GPNMB) emerged as one of the most strongly upregulated transcripts in DTPs, and also was amongst the genes associated with resistance in the PEARL dataset. We further verified that GPNMB overexpression (GPNMB-OE) in sensitive cells conferred resistance to CDK4/6 inhibition, and enhanced migratory capacity. Overexpression of GPNMB drove substantially faster tumor progression and eliminated the growth-inhibitory effect of abemaciclib, which remained highly effective in control tumors. Across all treatment arms, GPNMB-OE tumors failed to respond to CDK4/6 blockade, highlighting a strong resistance phenotype. These results identify GPNMB as a potent promoter of tumor progression and a key mediator of resistance to abemaciclib. Our findings position GPNMB as a potential biomarker and therapeutic target that may help identify patients unlikely to benefit from CDK4/6 inhibition.

## INTRODUCTION

Cyclin-dependent kinase (CDK) 4 and 6 inhibitors (CDK4/6i) have been one of the most significant practice-changing advancements in the treatment of estrogen receptor positive (ER+) breast cancer in the recent decade (1–3). CDK4/6i such as abemaciclib, palbociclib, and ribociclib have been approved for the treatment of metastatic ER+ breast cancer. In addition, abemaciclib and ribociclib are also US Food and Drug Administration (FDA)- and European Medicines Agency (EMA)-approved for treating early-stage breast cancer. CDK4/6i work by blocking the complex formation between CDK4/6 and cyclin D, which prevents the phosphorylation of the retinoblastoma protein (RB). This results in cell cycle arrest and prevents the uncontrolled proliferation (4).

Although many patients respond to CDK4/6i, some patients have intrinsic resistance to these agents while others may acquire resistance to therapy over time. Despite the initial response to CDK4/6i, resistance and recurrence are frequent contributing to breast cancer mortality. Most clinical trials with CDK4/6i in metastatic ER+ breast cancers do not require any biomarker analysis as entry criteria. The monarchE clinical trial in high-risk early breast cancer required Ki67 greater than 20% as a risk defining factors, which ultimately resulted in FDA-approval of Ki67 as a selection marker. However, responses to abemaciclib was observed in even in low-Ki67 patients, resulting in the removal of Ki67 as an FDA-approved biomarker for abemaciclib (5). The NATALEE clinical trial showed that adding ribociclib to endocrine therapy produced a meaningful and durable reduction in invasive disease-free recurrence across a broad group of intermediate- and high-risk HR+/HER2- early breast cancer (6, 7). This benefit was consistent across all clinicopathologic subgroups. However, no predictive biomarker has been identified to guide selection.

Several studies have analyzed samples from patients with metastatic breast cancer treated with CDK4/6i to identify the cause(s) of resistance. Genomic profiling has identified mutations in cell cycle-associated genes such retinoblastoma (RB) tumor suppressor in samples from patients treated with CDK4/6i (8). However, these were found to be rare (<5%) in patients enrolled in the PALOMA-3 clinical trial (9). Analysis of the monarchE and NATALEE studies did not find any alterations to be significantly associated with outcomes. Recent work by Kudo et al. has provided important genomic insights into resistance to first-line CDK4/6 inhibition in metastatic HR+/HER2– breast cancer (10). By analyzing MSKCC-IMPACT sequencing data from patients treated with CDK4/6i plus endocrine therapy, a striking enrichment of TP53 and MDM2 alterations was present in roughly 30% of early progressors with markedly shorter progression-free survival compared with long-term responders. These findings were subsequently validated in a large, pooled dataset of 4,457 patients from the MONALEESA and monarchE clinical trials, emphasizing the robustness of these genomic predictors. Complementary functional studies demonstrated that dual inhibition of CDK4/6 and CDK2 can mitigate the effects of p53 pathway disruption and restore durable therapeutic responses in ER+ breast cancer models, highlighting the biological relevance of p53 alterations in shaping CDK4/6i sensitivity and supporting rational combination strategies to overcome resistance.

Studies involving mRNA analysis of patient samples treated with CDK4/6i have largely failed to identify definitive markers associated with outcomes (11, 12). An exception to this was the analysis of the PEARL trial which identified the expression of a number of genes (41 refractory and 10 sensitive) including Cyclin E1 (*CCNE1*) and Polo-like kinase 1 (*PLK1*) as important determinants of resistance (13). Kong et al also developed a 33-gene signature that predicts neoadjuvant Ki67 response to anastrozole/palbociclib in a metastatic phase 2 trial (14). Other postulated mechanisms of CDK4/6i resistance in preclinical models include elevated CDK6 activity, cyclin E, Fibroblast Growth Factor Receptor (FGFR) pathway activation, pyruvate dehydrogenase kinase 1 (PDK1) and protein Arginine Methyltransferase 5 (PRMT5) (15–19). None of these molecular biomarkers are currently used for selecting patients for CDK4/6 inhibitors in clinical trials (20–23).

Close inspection of clinical trial data suggests two important facets associated with resistance to CDK4/6i: a) resistance to the drug can appear early with up to 25% of patients exhibiting disease progression within the first few months, b) Data from the MAINTAIN and postMONARCH clinical trials in metastatic BC showed that patients who had tumors that progressed on primarily palbociclib could be successfully treated with ribociclib or abemaciclib, respectively (24, 25). The transient nature of CDK4/6i resistance has been also observed in cell line-based studies (26). These studies suggest that mechanisms of resistance may be transient and perhaps the “drug-tolerant persisters” (DTPs) might provide a good model to study resistance. These cells are believed to be a subpopulation of slowly replicating cells that can tolerate the drug treatment and repopulate the tumor following the termination of therapeutics (27–29). DTPs have been implicated as a bridge for an early stage of drug resistance mechanism allowing the survival of cells and adaptation to secondary resistance phenotypes (reviewed by Pu et al. (30)). Although first described in lung cancer (31), DTP cells has been reported in multiple cancer types, including breast cancer (32–34).

In this study, we have developed abemaciclib and palbociclib resistant DTP models to study resistance to CDK4/6i. By comparing the DTP transcriptomic profiles with data from the PEARL clinical trial, we identified GPNMB, a transmembrane glycoprotein, as consistently overexpressed, supporting its role in resistance to CDK4/6i. Functional studies demonstrated that GPNMB promotes cell growth and contributes to CDK4/6i resistance. We further demonstrated its importance in tumor progression and resistance to abemaciclib. Together, these findings, suggest that GPNMB may serve as a promising biomarker for identifying patients unlikely to benefit from CDK4/6 inhibition.

## Materials and Methods

### Breast cancer cell lines and generation of DTPs

Human LCC2 (tamoxifen-resistant) and LCC9 (fulvestrant (ICI 182,780) and tamoxifen cross-resistant) cell lines were kind gifts from Dr. R. Clarke (Georgetown University Medical School, Washington DC)(35, 36). MCF7 (RRID: CVCL_0031), and T47D (RRID: CVCL_0553) cell lines were purchased from American Type Culture Collection (ATCC, Manassas, VA, USA). Cell lines were carefully maintained in a humidified tissue culture incubator at 37°C in a 5% CO2:95% air atmosphere, and stocks of the earliest passage cells were stored. Parental cells (LCC2, LCC9, MCF7 and T47D) that are sensitive to the relevant drugs (abemaciclib or palbociclib) were treated at concentrations exceeding 100 times the established IC_50_ values as described previously by Sharma et al. (31). DTPs appeared at 9 days later for the experimental analyses as indicated by Sharma et al. (31). In cell culture experiments, vehicle controls were used according to the solvents in which the drugs were prepared. Abemaciclib was dissolved in ethanol, whereas palbociclib was dissolved in dimethyl sulfoxide (DMSO). Accordingly, ethanol and DMSO were used as the respective vehicle controls at the same final concentrations as in the corresponding drug-treated groups (≤ 0.1% v/v).

### Cell proliferation

Cell viability was performed using the CyQuant Cell Proliferation Assay (#C35011) according to the manufacturer’s instructions (ThermoFisher Scientific, Waltham, MA, USA). Abemaciclib and palbociclib were purchased from Selleck Chemicals LLC (Houston, TX, USA). Briefly, cells were plated in 96-well flat-bottom plates at a density of 2,000 cells per well and allowed to attach overnight. The following day, cells were exposed to increasing concentrations (100–2000 nM) of abemaciclib or palbociclib for 72 hours. After treatment, CyQUANT reagent containing the fluorescent dye and membrane-permeabilizing agent was added according to the manufacturer’s instructions. Fluorescence was measured using a BioTek Synergy H1 plate reader (Agilent, Santa Clara, CA, USA) with excitation 480 nm and emission 535 nm. The relative fluorescence units (RFU) were used as an index of cell proliferation, and IC₅₀ values, the concentrations of inhibitors that are necessary to kill 50% of the cells, were calculated using GraphPad Prism 10.3.1.

### Cell cycle analysis by flow cytometry

Drug-sensitive cells and DTPs (2 x 10^5^) were plated on a 60-mm plate, harvested by trypsinization, pelleted, and resuspended in 1 mL of PBS. Cell cycle analysis was done as per manufacturer’s instruction (Propidium Iodide Flow Cytometry Kit, #ab139418, Abcam, Waltham, MA, USA) using a Becton Dickinson FACScan flow cytometer (Bedford, MA, USA). Data were analyzed with FlowJo™ v10.

### Senescence analysis

Senescence-associated β-galactosidase (SA-β-gal) expression was evaluated by using a Senescence β-Galactosidase Staining Kit (#9860, Cell Signaling Technology, Danvers, MA) and a flow cytometric detection of cellular senescence via β-galactosidase hydrolysis as per the manufacturer’s instructions (CellEvent Senescence Green Flow Cytometry Assay, #C10840, Thermofisher Scientific, Waltham, MA, USA). Flow cytometry analysis was performed using BD FACSuite software and Data were analyzed with FlowJo™ v10. DTPs or parental cells are quantified by the extent of senescent cells (green) to determine the senescent cells induced by abemaciclib or palbociclib.

### ALDEFLUOR assay

ALDEFLUOR assay was performed using the ALDEFLUOR Kit (StemCell Technologies, #01700) according to the manufacturer’s instructions. Briefly, cells were resuspended in ALDEFLUOR assay buffer containing the ALDH substrate (BAAA) and incubated at 37 °C for the recommended time, with a parallel sample treated with the specific ALDH inhibitor DEAB as a negative control. After incubation, cells were washed, kept on ice, and immediately analyzed by flow cytometry. ALDH-positive cells were identified by comparing fluorescence intensity of DTP and their sensitive counterparts to their DEAB-treated controls, and data were analyzed using BD FACSuite software and FlowJo™ v10.

### RNA isolation and sample quality assessment

Total RNA from DTPs (9 days treatment) and its parental drug-sensitive counterparts was extracted using the RNeasy isolation kit according to the manufacturer’s instructions (Qiagen, Germantown, MD, USA). RNA was treated with Turbo DNase (ThermoFisher Scientific, Waltham, MA, USA). The quality of RNA was assessed using the Nanodrop® ND-1000 spectrophotometer (ThermoFisher Scientific) and the Agilent 2100 Bioanalyzer (Agilent Technologies, Santa Clara, CA, USA).

### Clariom D Pico Human Transcriptome array and functional enrichment analysis

Total RNAs from DTPs and their parental drug-sensitive cell lines were sent to the Applied Biosystems/Thermo Fisher Scientific Service laboratory (Santa Clara, CA, USA). Quality Control and Clariom D Pico Human Transcriptome Array were performed according to Applied Biosystems/Thermo Fisher Scientific’s instructions as previously described (37). Probe cell intensity (CEL) files generated from Clariom D arrays were analyzed using Transcriptome Analysis Console (TAC) software version 4.0 (Applied Biosystems/ThermoFisher). Differential gene expression analysis was performed using a limma-based linear modeling framework with empirical Bayes–moderated statistics. P values and log₂-based fold changes were calculated for each comparison, and statistically significant genes were identified based on predefined thresholds. Differentially regulated genes (DEGs) were categorized as upregulated and downregulated sets in DTPs versus parental sensitive counterparts for each cell line. Venn diagrams were generated using genes that met the significance criteria of |Fold Change| ≥ 2 and *P* < 0.05, using the Venn diagram tool from Ghent University. of Ghent https://bioinformatics.psb.ugent.be/webtools/Venn/. Pathway enrichment analysis was conducted using g:Profiler (https://biit.cs.ut.ee/gprofiler/). Enriched pathways were identified across Gene Ontology (GO Biological Process).

### Quantitative reverse transcription-polymerase chain reaction (RT-qPCR) of breast cancer cell lines

RT-qPCR from DTPs and their parental drug-sensitive counterparts was performed as described previously (37). The mRNA levels of *GPNMB* (Hs01095669_m1) were analyzed using TaqMan gene expression assays on an ABI Prism 7900 platform according to the manufacturer’s instructions (Applied Biosystems/ThermoFisher Scientific, Carlsbad, CA, USA). Actin (*ACTB*; Hs00357333_g1) was used as endogenous control. All experiments were performed as three independent sets. The relative quantification of the gene expression changes was analyzed according to the ΔΔCt method using the Applied Biosystems DataAssistTM Software v3.0. All graphs were generated using GraphPad Prism 10.3.1 software. The error bars were calculated and represented in terms of the mean ± SD. The results presented are the combination of three independent assays, and unpaired t-test with Welch’s correction analyses were performed using GraphPad Prism 10.3.1 software (P < 0.05, statistically significant).

### Cell Migration Assay (Wound Healing Assay)

To assess the effects of GPNMB, 24 well plates were coated with either 0.1% bovine serum albumin (BSA) or recombinant human GPNMB (osteoactivin (rhOA; 10 μg/mL, Acro Biosystems, # catalog GPB-H5229). Briefly, 250 μL of freshly prepared coating solution was added per well and incubated at 37°C for 2 hours or at 4°C overnight. Wells were then aspirated, washed once with 1× PBS, and used immediately or within one week. Wound healing assay has been performed to measure two-dimensional migration (34). Briefly, cells were seeded at a density of 2 × 10^5 cells/mL in coated 24-well plates and cultured for 72 hours to allow monolayer formation. A linear scratch was generated using a p200 pipette tip, and detached cells were removed by gentle washing with PBS. Images were acquired at the indicated time point using a Nikon Eclipse TS100 microscope with NIS-Elements imaging software (version 5.41.02; Nikon Corporation). Wound area was quantified, and percent wound closure was calculated using the formula: [(wound area at time 0 − wound area at time x) / wound area at time 0] × 100. Data were analyzed and graphed using GraphPad Prism v10.3.1 software. Results represent three independent experiments performed in triplicate. Statistical analyses were conducted using an unpaired two-tailed Student t test with Welch’s correction. Differences were considered statistically significant at P < 0.05.

### Western Blot

The protein lysates from DTPs and their parental drug-sensitive counterparts were prepared, and equal amounts of protein were subjected to SDS–PAGE and Western blot analysis as described previously (38). The Bio-Rad DC-Protein assay kit (Bio-Rad, Hercules, CA, USA) was used to determine protein concentrations. Blots were incubated with the primary antibody against GPNMB (Cat# 38313 Cell Signaling Technology, Danvers, MA). Antibody against β-actin (ACTB, Sigma, St. Louis, MO, USA, dilution 1:5000) or GAPDH (Cat# GTX627408, GeneTex, Inc., Irvine, CA, USA) was used as the loading control. Protein bands were visualized by SuperSignal West Pico PLUS Chemiluminescent Substrate kit (Amersham, Piscataway, NJ, USA) and Amersham Imager 600 GE Healthcare Life Sciences (GE Healthcare Bio Sciences, Pittsburgh, PA, USA). The data are representative of three individual sets.

### Spheroid Immunofluorescence

Spheroid development and immunofluorescence staining were performed as described previously (39). Briefly, primary antibody staining was performed using GPNMB (Cat# sc-271415, Santa Cruz, CA, USA, dilution 1:50). Secondary antibody F(ab’)2-Goat anti-Mouse IgG (H+L) Cross-Absorbed, Alexa Fluor Plus 555 was purchased from Invitrogen, cat#A48287, dilution 1:1000). Spheroids were imaged with the Leica TCS SP8 inverted confocal microscope (X 20). DAPI for nuclear staining is purchased by Cell Signaling (Cat#4083).

### Generation of stable full-length GPNMB overexpression in breast cancer cells

Lentiviral particles encoding the human *GPNMB* ORF (NM_001005340, GPNMB Human Tagged Lenti ORF clone, (OriGene cat# RC207615L4V, pLenti-C-mGFP-P2A-Puro) were obtained from OriGEne Technologies (Rockville, MD, USA). The pLenti-C-mGFP-P2A-Puro vector (OriGene cat# PS100093V) served as the lentiviral control for all experiments and labeled as empty vector (EV). LCC2, MCF7 and T47D cells were transduced with either GPNMB ORF (GPNMB Overexpression, GPNMB-OE) or control lentivirus (Empty Vector-EV) according to the manufacturer’s instructions. After transduction, cells were selected with puromycin to establish stable cell lines expressing GPNMB or vector control. Individual puromycin-resistant clones were isolated, expanded, and screened for GPNMB expression by Western blotting and GFP fluorescence. The clone demonstrating the highest GPNMB overexpression (GPNMB OE clone 1) from each cell line model was selected for all subsequent assays. mRNA and protein expression levels of GPNMB were further validated using RT-qPCR and immunofluorescence, respectively.

## Statistical analysis

Unpaired t-test with Welch’s correction analyses were performed using GraphPad Prism 10.3.1 software as indicated. All results are representative of three replicates and expressed as mean values SD. In all cases, differences were considered statistically significant at \**P* < 0.05, \*\**P*<0.01, \*\*\**P*<0.001, \*\*\*\**P*<0.0001.

### *In vivo* orthotopic xenograft model

All animal experiments were conducted under a protocol approved by the Emory University Institutional Animal Care and Use Committee (IACUC). Six-to eight-week-old ovariectomized athymic nude mice (Jackson Lab, Bar Harbor, ME, USA) were acclimatized for 3–7 days. All animals were housed in a specific pathogen-free (SPF) facility at Emory University. A controlled-release E2 pellet (0.72 mg E2, 60-day formulation; Innovative Research of America, Sarasota, FL, USA) was injected subcutaneously (s.c.) via a sterile 14-gauge trocar 24 h before tumor implantation. LCC2 cells stably expressing either empty vector (2EV) or GPNMB (2GPNMB OE C1) were injected into the mammary fat pad at a density of 5 × 10⁶ cells per mouse. Tumor growth was monitored by caliper measurements, and tumor volume was calculated using the formula: (length × width²)/2. When tumors reached approximately 100–150 mm³ (day 24 post-injection), mice were randomized into four treatment groups: (1) LCC2-EV (2EV) treated with vehicle (n = 9), (2) LCC2-GPNMB (2GPNMB OE C1) treated with vehicle (n = 9), (3) LCC2-EV (2EV) treated with abemaciclib (n = 10), and (4) LCC2-GPNMB (2GPNMB OE C1) treated with abemaciclib (n = 10). Abemaciclib (Selleckchem, Cat # S5716) was formulated as a suspension in 0.5% methylcellulose for delivery via oral gavage (PO) at a dose of 50 mg/kg/once daily for 35 consecutive days. Vehicle only (0.5% methylcellulose) - treated mice received an equivalent volume by oral gavage. Animals were sacrificed after 35 days of treatment when tumors of 2GPNMB OE C1 (the group with the highest tumor growth) reached a volume of 1,000 mm^3^.

### Statistical analysis *in vivo*

Tumor growth data are presented as mean ± SEM. Statistical significance between treatment groups was determined using two-way ANOVA with repeated measures, followed by appropriate post hoc multiple-comparison tests. An adjusted (adj.) *p* value < 0.05 was considered statistically significant. All statistical analyses were performed using GraphPad Prism v10.3.1 software.

## Results

### Drug -Tolerant Persisters (DTPs) to CDK4/6i display distinct functional phenotypes compared to their drug naive sensitive counterparts

To investigate CDK4/6i resistance, we developed DTP cell models resistant to CDK4/6i (abemaciclib or palbociclib) in a panel of ER+ cells (LCC2 control, LCC9 control, MCF7 control and T47D control) treated as 100x IC_50_ doses. This resulted in a subpopulation of cells resistant to CDK4/6i (i.e., DTPs) after 9 days of treatment (**Fig. 1A**). The DTPs failed to reach 50% inhibition at the highest concentrations tested (2000nM), indicating that the cells became resistant to both abemaciclib and palbociclib. LCC2 control, sensitive cells to abemaciclib and palbociclib, exhibited a potent dose-response with an IC₅₀ of 47.32 nM and 49.96 nM, respectively, as described previously (40). A similar pattern was observed in the other DTP sublines confirming that this is general process ((40) and **Supplementary Fig. S1A**). The Palbociclib-DTPs similarly failed to reach 50% growth inhibition across the tested concentration range, further confirming resistance ((40) and **Supplementary Fig. S1A**).

**Fig 1.**
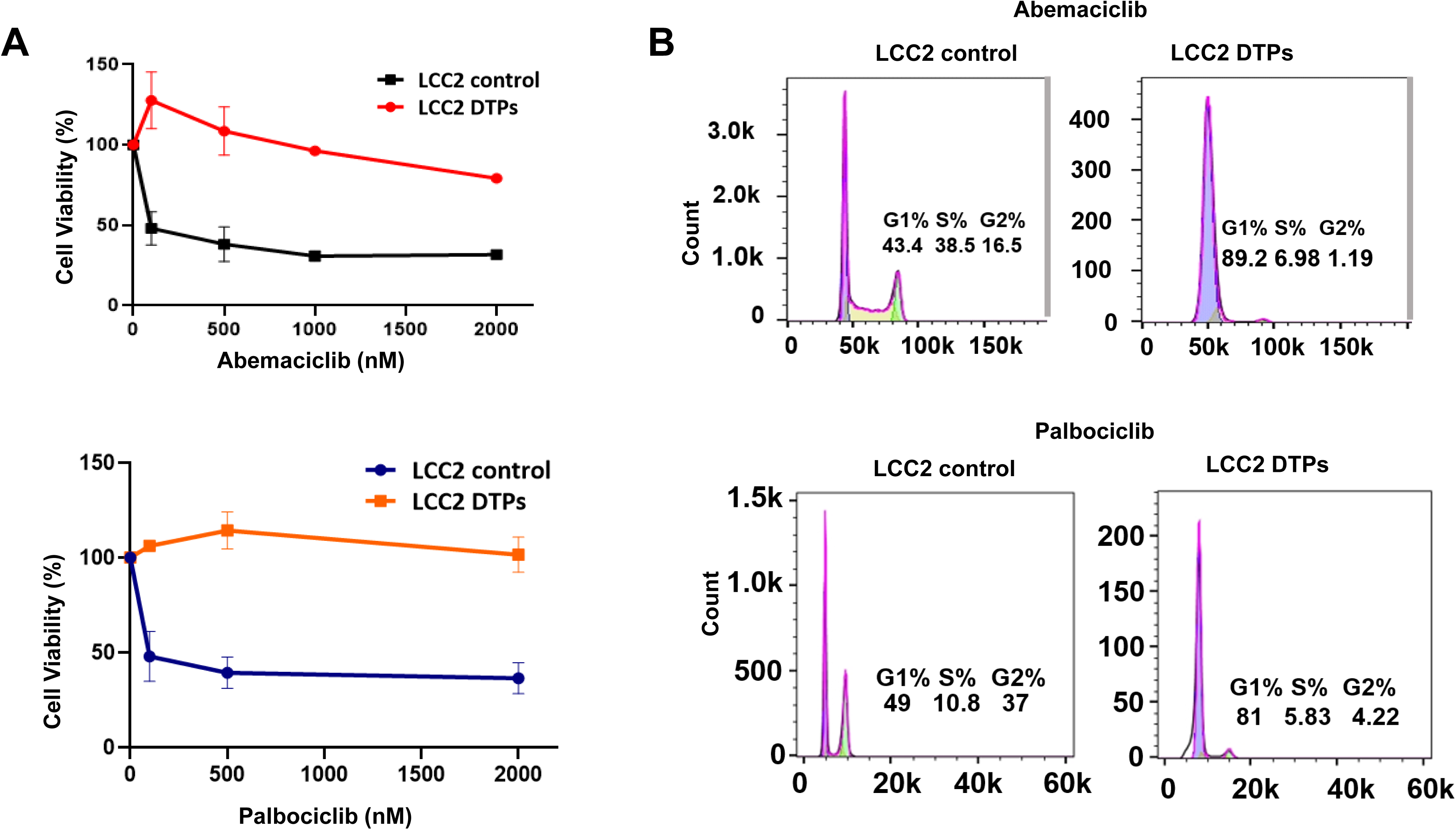
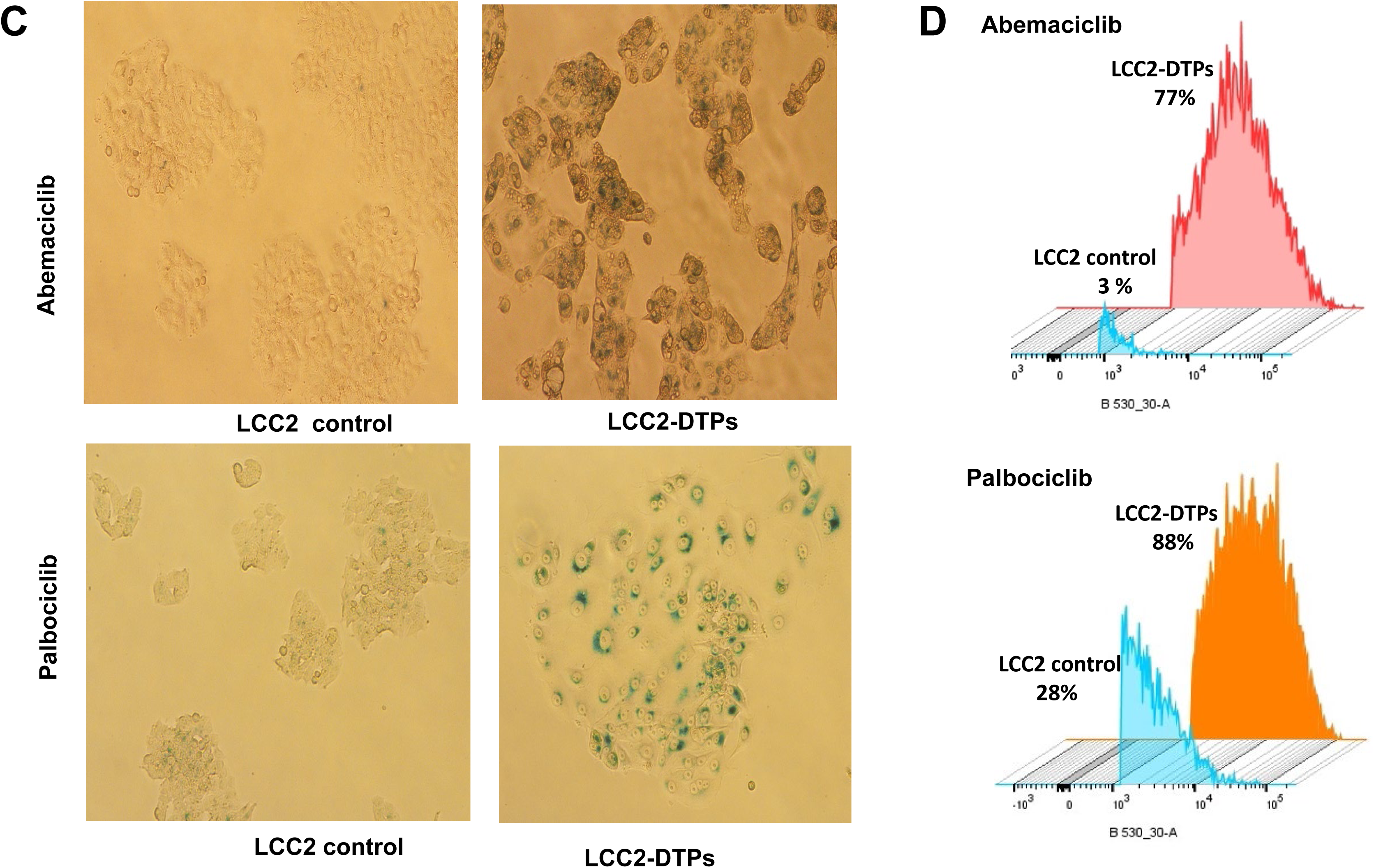
Phenotype of CDK4/6i-treated drug tolerant persister cells (LCC2 DTPs) versus sensitive (LCC2 control) cells. **A** Representative sensitivity of LCC2 control treated with vehicle and LCC2 DTPs treated with abemaciclib (top) and palbociclib (bottom) with the indicated concentrations for 72hr using CyQUANT Direct Cell Proliferation Assay. IC50 Data (n = 3) were calculated using GraphPad Prism. **B** Representative cell cycle distribution of the LCC2 control and LCC2-DTPs developed to abemaciclib (top) and palbociclib (bottom). Cells at G1, S, and G2-M phases for each condition are shown in percentage (mean ±SD, n = 3). **C** Representative senescence-associated β-galactosidase (SA-β-gal) expression using Senescence β-Galactosidase Staining Kit and **D** Representative FACS plots showing SA-β-gal expression in LCC2 control and LCC2 DTPs developed to abemaciclib (top) and palbociclib (bottom) using CellEvent Senescence Green probe specific to β-gal. Numbers indicate the percentage of senescent cells.

To determine whether the DTP phenotype altered the canonical G1-arrest response associated with CDK4/6 inhibition, we next evaluated cell-cycle distribution following treatment with abemaciclib and palbociclib. As expected, LCC2, LCC9, MCF-7, and T47D DTP sublines showed G1-phase growth arrest upon exposure to both inhibitors (**Fig. 1B, Supplementary Fig. S1B)**. Because prolonged or repeated G1 arrest induced by CDK4/6 inhibition can drive cells toward a senescent state, we next asked whether DTPs exhibit enhanced senescence compared with their sensitive parental counterparts. We previously demonstrated that acquired resistant cells to abemaciclib or palbociclib developed a senescent phenotype (40). To determine whether drug-tolerant persisters (DTPs) similarly displayed increased senescence, we measured senescence-associated β-galactosidase (SA-β-Gal) activity using histochemical staining and flow cytometry, a widely accepted surrogate marker of cellular senescence. In LCC2 DTP sublines, we observed marked blue staining, indicative of elevated SA-β-Gal activity and enhanced senescence (**Fig. 1C**). Quantification of mean fluorescence intensities further confirmed these findings: LCC2 DTPs generated with abemaciclib showed a 77% increase in SA-β-Gal–positive cells, while LCC2 DTPs generated with palbociclib exhibited 88% positivity, compared with only 3% and 28% in the respective parental controls (p < 0.0001) (**Fig. 1D**)). Similar results were observed in additional DTP models (LCC9, MCF7, and T47D), emphasizing that senescence is a prominent characteristic of CDK4/6 inhibitor-induced drug tolerance (**Supplementary Fig. S1C-D**).

It has been reported that ALDH activity, particularly ALDH1, is essential for maintaining a subpopulation of DTP cells that survive prolonged exposure to targeted therapies (41). We next examined whether ALDH1 contributes to the drug-tolerant persister (DTP) phenotype to abemaciclib and palbociclib. Using the ALDEFLUOR assay, we did not detect any measurable ALDH1 activity in DTP cells (**Supplementary Fig. S2**). This absence of ALDH1 activity argues against a stem-cell–like mechanism driving persistence in our model. Moreover, DTPs fully reverted to the parental drug-sensitive phenotype after 20 days in drug-free media, consistent with the transient and reversible nature of persister states reported in clinical settings (**Supplementary Fig. S3**).

### Transcriptomic analysis in DTPs

We next identified differentially expressed genes (DEGs) in four DTP cell lines treated with abemaciclib compared to their drug naïve sensitive counterparts using the Clariom D transcriptomics arrays and TAC 4.0 platform (*Set 1-LCC2-DTPs versus LCC2 control, set 2-LCC9 DTPs versus LCC9 control, set 3-MCF7 DTPs versus MCF7 control and set 4- T47D DTPs versus T47D control)*. This analysis revealed the upregulation of 154 genes common in all DTP models, while 274 common downregulated genes were identified (**Fig. 2A**). To gain deeper insight into the molecular programs altered in DTPs, we performed pathway enrichment analysis using g:Profiler with GO Biological Process annotations. Consistent with the known mechanism of CDK4/6 inhibition, the most significantly downregulated genes were predominantly enriched in pathways related to cell cycle progression and cell division, confirming the suppression of proliferative programs. In contrast, the top upregulated genes were significantly associated with biological processes including response to growth factor, cell migration, and integrin-mediated signaling pathways (**Fig. 2B).** Together, these findings indicate that while CDK4/6i primarily suppresses proliferative programs, DTPs simultaneously activate signaling networks that may facilitate persistence and cellular remodeling.

**Fig. 2.**
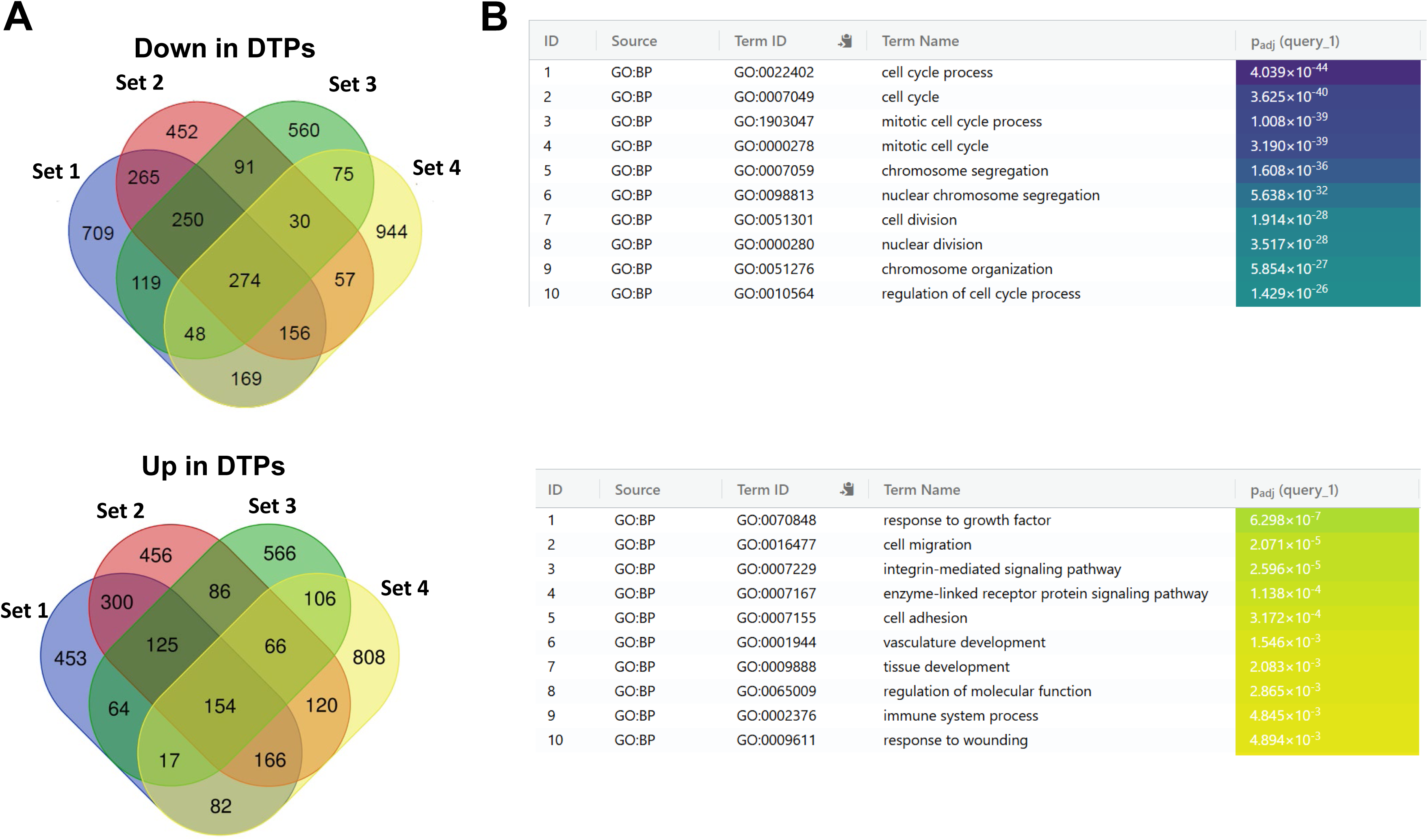
Transcriptomic analysis of DTPs to CDK4/6i reveals enrichment of ECM pathways in comparison to their sensitive counterparts. **A** Venn diagram showing the differential expression analysis of DTPs compared with cells with their drug sensitive counterparts calculated by Transcriptome Analysis Console (TAC 4.0) software (Top: downregulated genes in DTPs, 2 < fold change > 2 cutoff (P < 0.05); bottom: upregulated genes in DTPs): *Set 1-* LCC2 DTPs versus LCC2 control*, set 2-* LCC9 DTPs versus LCC9 control, *set 3*- MCF7 DTPs versus MCF7 control and *set 4*- T47D DTPs versus T47D control. B g:profiler pathway analysis for down-regulated genes (top) and up regulated genes (bottom) in DTPs compared their sensitive counterparts.

### GPNMB Emerges as a Top Differentially Expressed Gene Associated with CDK4/6 Resistance in ER+ Breast Cancer

To identify a clinically relevant dataset for validating the 154 genes upregulated in DTPs, we systematically searched for clinical trials that included both gene-expression profiling and treatment with CDK4/6 inhibitors. This search identified the phase III PEARL trial, which randomized participants to palbociclib plus endocrine therapy versus capecitabine, as the only publicly available cohort that combined large-scale mRNA expression data with detailed clinical outcomes from patients with HR+/HER2- metastatic breast cancer treated with palbociclib plus endocrine therapy (13). Among the forty-one refractory genes previously linked to resistance to palbociclib plus endocrine therapy (13), glycoprotein non-metastatic B (GPNMB) emerged as one of the most strongly upregulated transcripts in our DTP models [FDR at 5% with the q-value method (SAM)], further supporting its role as a clinically validated marker of CDK4/6 inhibitor resistance.

### *GPNMB* is significantly elevated in CDK4/6i-DTPs and overexpression of GPNMB in vitro confers resistance

High expression of *GPNMB* mRNA was verified in LCC2-derived DTP sublines to abemaciclib (100-fold; *P*<0.0038) and palbociclib (33-fold; *P*<0.0042) compared to their LCC2 control cells (**Fig. 3A).** This significant upregulation was further verified in additional ER-positive, DTP sublines (LCC9 DTPs, MCF7 DTPs and T47D DTPs) in both abemaciclib and palbociclib, demonstrating that GPNMB upregulation is not restricted to a single cell-line model **(Supplementary Fig. S4A-B).** Flow cytometry analysis also verified that DTPs generated in response to abemaciclib or palbociclib exhibited a dramatic elevation of GPNMB at the cell surface, indicating strong upregulation of this protein during CDK4/6 inhibition (**Fig. 3B; Supplementary Fig. 5).** Using spheroid immunofluorescence, we further confirmed the upregulation of GPNMB at the protein level in all DTP sublines (**Fig. 3C; Supplementary Fig. 6**).

**Fig. 3.**
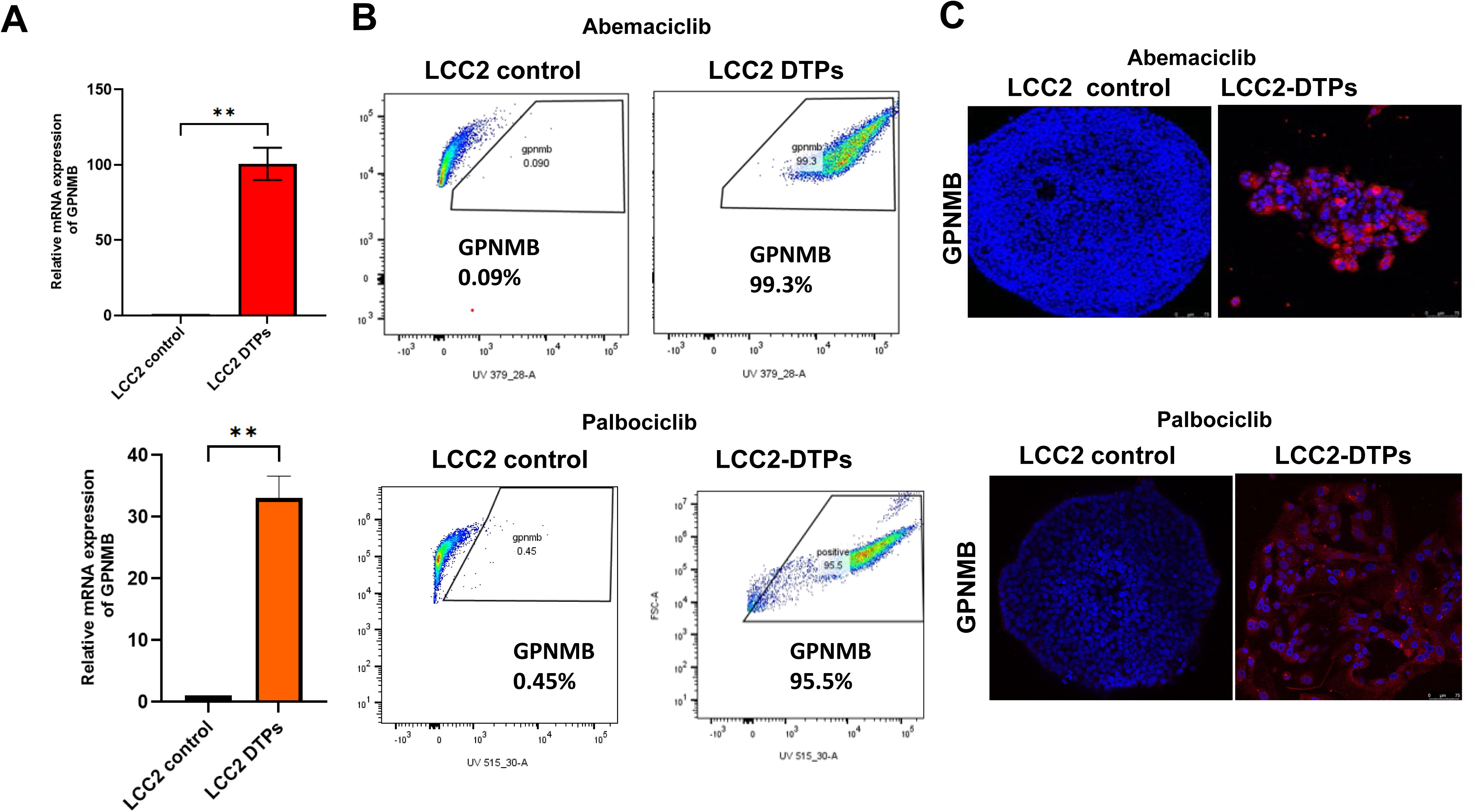
GPNMB is elevated in CDK4/6i-DTP cells at both mRNA and protein levels. **A** Fold change (mean ± SD, n = 3; \*\**P* < 0.01) of GPNMB mRNA expression levels were assessed by RT-qPCR in CDK4/6i-tolerant persisters (LCC2-DTPs) compared with LCC2 control cells. Unpaired t-test with Welch’s correction analyses were performed using GraphPad Prism 10.3.1 software as indicated **B** FACS-based analysis of cell-surface GPNMB expression in LCC2-DTPs compared to LCC2 control cells. High GPNMB cell surface expression in DTPs with the GPNMB antibody. **C** Fluorescent imaging of GPNMB protein in LCC2 DTPs and LCC2 control. GPNMB (Cat# sc-271415, Santa Cruz, CA, USA, dilution 1:50). Secondary antibody F(ab’)2-Goat anti-Mouse IgG (H+L) Cross-Absorbed, Alexa Fluor Plus 555)

### Overexpression of GPNMB in parental sensitive cells mimic the DTP phenotype

To further investigate the role of GPNMB in conferring resistance to CDK4/6 inhibitors, we generated stable GPNMB-overexpressing LCC2 parental breast cancer cells using the OriGene human GPNMB ORF construct, followed by antibiotic selection and screening of individual clones. The clone with the highest GPNMB expression in LCC2 (named GPNMB clone 1; 2GPNMB-OE-C1) was selected for further functional studies (**Supplementary Fig. S7A)**. We confirmed the dramatic increase of *GPNMB* expression at mRNA levels (P=0.0002; **Fig. 4A**). We also confirmed the elevated increase of GPNMB at the protein levels (**Fig 4B)**; We next determined the sensitivity of GPNMB-overexpressing cells to CDK4/6i (abemaciclib). 2EV control cells displayed an IC₅₀ of approximately 50 nM, whereas 2GPNMB-OE-C1 cells exhibited a dramatically elevated IC₅₀ of 1047 nM (**Fig. 4C**). This shift represents nearly a 20-fold increase in resistance relative to parental cells, indicating that GPNMB overexpression substantially diminishes the efficacy of CDK4/6 inhibition. Consistent with its impact on cell-cycle regulation, 2GPNMB-OE-C1 resulted in an elevated accumulation of cells in the G1 phase (**Supplementary Fig. S7B**). This G1 arrest was accompanied by a corresponding increase in senescence (**Supplementary Fig. S7C)**, further supporting the role of GPNMB in modulating proliferative capacity and cell-cycle progression. Together, these findings demonstrate that GPNMB overexpression not only alters cell-cycle dynamics but also confers resistance to abemaciclib, highlighting a potential mechanism by which tumor cells may evade CDK4/6-targeted therapy.

**Fig. 4.**
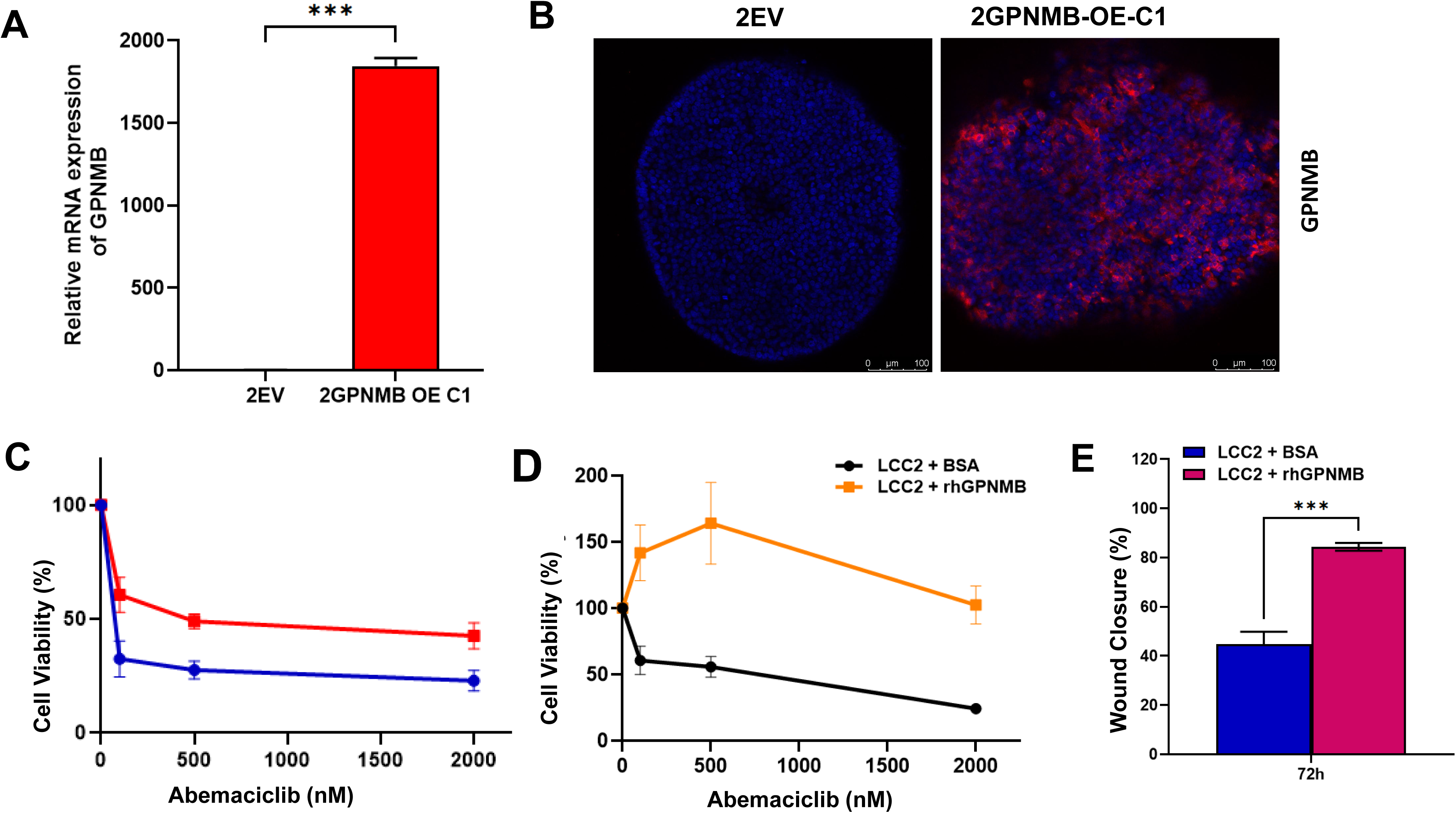
Expression analysis of GPNMB overexpression in LCC2 cells. LCC2 cells were transduced either with empty vector (2EV) or GPNMB construct (2GPNMB-OE-C1) using lentiviral based transduction system. **A** GPNMB mRNA verification in 2GPNMB-OE-C1 compared to 2EV using RT-qPCR. Unpaired t-test with Welch’s correction analyses were performed using GraphPad Prism 10.3.1 software (\*\*\**P*<0.001). **B** GPNMB protein expression using spheroid immunofluorescent assay. **C** Representative sensitivity of 2EV and 2GPNMB-OE-C1 cells treated with the indicated concentrations of abemaciclib for 72hr using CyQUANT Direct Cell Proliferation Assay (n = 3). **D** Representative sensitivity to abemaciclib and **E** percentage of wound closure coverage healing were performed using LCC2 cells plated on rhGPNMB or 0.1 % BSA coated dishes; (n = 3). For statistical significance, unpaired t-test with Welch’s correction analyses were performed using GraphPad Prism 10.3.1.

**Fig. 5.**
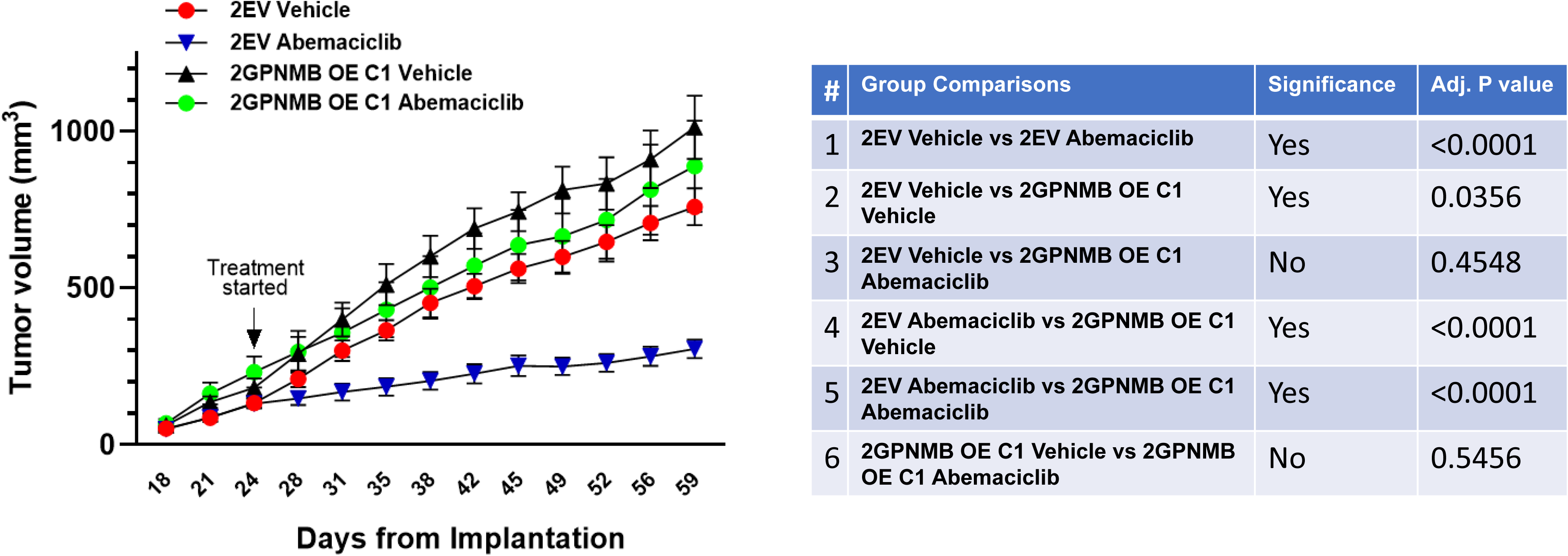
Overexpression of GPNMB results in significant tumor growth and confers resistance to abemaciclib. Tumor growth and response to abemaciclib were assessed in mice bearing empty vector (2EV) or GPNMB overexpression (2GPNMB OE C1) in LCC2 mammary xenografts and treated with vehicle as a suspension in 0.5% methylcellulose or abemaciclib at a dose of 50 mg/kg/once daily for 35 consecutive days via oral gavage (PO). Four experimental groups were analyzed: (1) 2EV Vehicle (red symbol), (2) 2EV Abemaciclib (blue symbol), (3) 2GPNMB OE C1 Vehicle (black symbol), and (4) 2GPNMB OE C1 Abemaciclib (green symbol). Tumor growth data are presented as mean ± SEM. Statistical significance between treatment groups was determined using two-way ANOVA with repeated measures, followed by appropriate post hoc multiple-comparison tests. An adjusted (adj.) p value < 0.05 was considered statistically significant. All statistical analyses were performed using GraphPad Prism v10.3.1 software.

We next confirmed that GPNMB conferred resistant to abemaciclib by culturing sensitive parental LCC2 cells on recombinant GPNMB protein. While cells coated with GPNMB protein did not reach IC50 values, sensitive control LCC2 cells coated with 0.1% BSA reached IC50 values of (439.2 nM, **Fig. 4D**). We next demonstrated its role in cell migration using a wound-healing assay performed on surfaces coated with either BSA or recombinant human GPNMB (GPNMB-ECD). LCC2 cells cultured on GPNMB-ECD–coated plates exhibited significantly enhanced wound closure compared with cells plated on BSA controls, indicating that extracellular GPNMB directly promotes migratory capacity (**Fig. 4E**; *P*=0.0003). Consistent with this observation, the addition of soluble recombinant GPNMB-ECD to parental LCC2 cells similarly accelerated wound closure, further supporting a functional role for the extracellular domain in driving a pro-migratory phenotype. These findings align with our resistance model, in which GPNMB upregulation is associated with reduced sensitivity to CDK4/6 inhibition. Together, these results demonstrate that the GPNMB extracellular domain is sufficient to enhance cell migration and contributes to the invasive and drug-resistant behavior observed in GPNMB-high breast cancer cells.

### Overexpression of GPNMB Promotes Tumor Growth and Abemaciclib Resistance in ERpositive breast cancer xenografts

To determine how GPNMB overexpression influences tumor growth and therapeutic response to abemaciclib, we evaluated four xenograft conditions: These included LCC2 empty-vector tumors treated with vehicle (2EV vehicle) and with abemaciclib (2EV abemaciclib), as well as LCC2 tumors overexpressing GPNMB (GPNMB-OE-C1) treated with either vehicle or abemaciclib. This design allowed us to directly compare the impact of GPNMB overexpression under both untreated and drug-treated conditions. Tumor growth in 2EV group was suppressed significantly in response to abemaciclib, confirming the sensitivity of 2EV group to abemaciclib (adjusted P<0.0001). On the other hand, GPNMB overexpression alone markedly enhanced tumor growth relative to 2EV tumors, demonstrating a significant oncogenic role for GPNMB *in vivo* (adjusted P<0.0356). Tumors expressing GPNMB (GPNMB-OE-C1) continued to grow and remained unresponsive to abemaciclib compared with the 2EV group, demonstrating that GPNMB overexpression confer resistance to abemaciclib. Direct comparisons further highlighted the resistance phenotype driven by GPNMB. 2EV tumors treated with abemaciclib were significantly smaller than GPNMB-OE-C1 tumors receiving either vehicle or abemaciclib, demonstrating that tumors with GPNMB overexpression acquire a phenotype that confers resistance to abemaciclib blockade (adjusted P<0.0001). When comparing GPNMB-OE-C1 tumors treated with abemaciclib to those receiving vehicle, tumor growth remained essentially unchanged, and the difference was not statistically significant, emphasizing the diminished therapeutic effect of abemaciclib in the context of GPNMB overexpression. These findings suggests that GPNMB overexpression not only drives aggressive tumor progression, but also confers resistance to abemaciclib, positioning GPNMB as a critical determinant of reduced responsiveness to abemaciclib.

## DISCUSSION

The transient nature of CDK4/6i resistance observed in the MAINTAIN (24) and postMONARCH (25) clinical trials supports the application of DTPs-based models for investigation of CDK4/6i resistance. DTPs have been shown to repopulate drug-sensitive cellular progenies following a ‘drug holiday’ without harboring any known resistance-associated secondary mutations (42). We used high-dose treatment of human BC cell lines with CDK4/6i (abemaciclib and palbociclib) to develop an effective model of DTPs-model to explore the molecular mechanisms associated with resistance. CDK4/6i-DTPs exhibited slow growth, senescence, and resistant to CDK4/6i.

We investigated the characteristics of CKD4/6i-DTPs to further understand these clinical observations. Transcriptomics analysis identified 154 upregulated transcripts (138 protein coding) to be differentially expressed in CKD4/6i-DTPs as compared to parental cells across the four cell lines analyzed. To assess the clinical relevance of these genes, we compared this data with genomics analysis of 2,597 genes in the phase III PEARL study (13), wherein patients were treated with palbociclib plus ET. GPNMB was the only gene identified among the 41 top differentially expressed genes (FDR at 5% with the q-value method (SAM)) associated with resistance to palbociclib and endocrine therapy. This provided clinical evidence supporting the importance of GPNMB in palbociclib resistance. In our models, we observed that most of the genes related to cell cycle and proliferation were downregulated in DTPs including CCNE1 and PLK1.

GPNMB is expressed by senescent cells as has been referred to as a seno-antigen. Cellular senescence is one of new hallmarks of cancer mediating tumor development and malignant progression. (43) Therapy-induced senescence, a cellular state triggered by therapeutics, has been implicated as a mechanism of resistance (44). In particular, reversible senescent cells such as in DTPs can escape from their non-proliferative condition and resume cell proliferation and rise to therapy-resistance and tumor progression. (45). In spite of this, the cells can show (near-)complete recovery to parental phenotype suggesting that they do not undergo “irreversible” senescence. Kudo et al have suggested that p53 deficiency suppresses the DREAM complex in breast cancer cells, which enables cell-cycle re-entry (46). We recently have shown that palbociclib- and abemaciclib-acquired resistant cells induce senescence and exhibit markers of therapy-induced senescence (40). However, we did not observe any change in expression of TP53 mRNA in both acquired resistance and DTP models (Supplementary Fig. S8). and CDK2 mRNA levels were down in DTPs, but not significant, suggesting that therapy-induced senescence and these processes are likely independent of TP53 signaling. Overexpression of GPNMB, a transmembrane protein, has been associated with therapeutic resistance and poor overall survival in several tumors, including breast and head and neck cancers (47–49). A role of GPNMB has been reported in the development of resistance to chemotherapy and PD-L1-directed immunotherapy (50, 51). Our study has identified that GPNMB-overexpression in breast cancer cells was associated with CDK4/6i resistance. Our *in vivo* studies in LCC2 xenograft model demonstrate that GPNMB overexpression profoundly alters both tumor growth dynamics and therapeutic responsiveness to the CDK4/6 inhibitor abemaciclib. In line with expectations, control (2EV) tumors exhibited a robust reduction in tumor growth following treatment with abemaciclib, confirming the inherent sensitivity of this model to CDK4/6 blockade. This finding reinforces the established role of abemaciclib as an effective inhibitor of cell-cycle progression in hormone-resistant breast cancer models.

In contrast, GPNMB overexpression significantly accelerated tumor growth even in the absence of treatment, emphasizing the potent oncogenic capacity of GPNMB *in vivo*. In addition, GPNMB overexpression not only promoted tumor growth but also markedly diminished the therapeutic efficacy of abemaciclib. Tumors with GPNMB overexpression were entirely unresponsive to abemaciclib, maintaining growth rates indistinguishable from their vehicle-treated counterpart. These results position GPNMB as a critical determinant of therapeutic resistance to CDK4/6 inhibition. The dual impact of GPNMB as enhancing tumor promotion while simultaneously reducing drug responsiveness highlights its relevance as both a prognostic marker and a potential therapeutic target. Strategies aimed at targeting GPNMB may restore sensitivity when combined with CDK4/6 inhibitors and improve outcomes in tumors with high GPNMB expression.

Therapeutic targeting of GPNMB has been attempted to counter aging and to autoimmune diseases (52, 53) in addition to cancer. Elimination of GPNMB-positive cells by senolytic GPNMB vaccination reversed pathological aging in aged mice and prolonged the lifespan of mice with premature aging. GPNMB has been well documented to be an important target in metastatic Triple Negative Breast Cancer (TNBC)(47). In TNBC, an antibody-drug conjugate, Glembatumumab vedotin (GV) has been studied (49). The phase II METRIC clinical showed that although GV had an acceptable safety profile, it was not associated with improvement in PFS as compared to capecitabine. The lack of clinical benefit in this trial has been attributed, at least in part, to GPNMB being predominantly in the cytosol in TNBC (49, 54) in contrast to our findings of cell surface GPNMB in CDK4/6i-DTPs and GPNMB-OE cells.

In summary, our study identifies GPNMB as a previously unrecognized driver of resistance to CDK4/6 inhibition. We demonstrate for the first time that GPNMB is markedly overexpressed at both the mRNA and protein levels in abemaciclib and palbociclib DTP models, and that its elevated expression is closely associated with an enhanced senescent phenotype as well as conferring resistance to CDK4/6i. These findings position GPNMB not only as a functional contributor to CDK4/6 inhibitor resistance, but also as a potential biomarker for stratifying patients who are more likely to experience reduced progression free survival on CDK4/6i therapy. Together, our results highlight GPNMB as a promising therapeutic and prognostic target that warrants further investigation in the context of overcoming or preventing CDK4/6 inhibitor resistance.

## Data Availability Statement

The CEL files for Clariom D Human Array will be deposited at GEO when the manuscript is accepted.

## Acknowledgements

This work was supported partly by the Susan G. Komen for the Cure to Sunil Badve (SAC220219) and Winship Invest$$ pilot grant to Yesim Gokmen-Polar. In addition, Startup funds from Emory University were also utilized. Sunil Badve is a Komen scholar. Research reported in this publication was supported in part by the Pediatrics/Winship Flow Cytometry Core of Winship Cancer Institute of Emory University, Children’s Healthcare of Atlanta (NIH/NCI) and the Cancer Animal Models shared resource of Winship Cancer Institute of Emory University under award number P30CA138292), and Emory University Integrated Cellular Imaging Core and Children’s Healthcare of Atlanta, (RRID:SCR_023534). The content is solely the responsibility of the authors and does not necessarily reflect the official views of the National Institute of Health.

## Ethics Approval / Consent to Participate

All experimental procedures were approved by the Emory University Institutional Biosafety Committees. All protocols related to patient samples were reviewed and approved by the Institutional Review Board (IRB) of Emory University. Samples and clinical records were de-identified prior to access by the authors and linked with a numerical identifier. The IRB waived requirement of informed consent.

## Author Contributions

YG-P, and SSB developed the study concept and design, obtained funding, and performed the interpretation, manuscript writing and execution of the entire project; YG and LR performed the experiments and statistical analyses. YH and MG-R carried out in vivo experiments and analyses, TB guided the entire project, KMK advised the conduct of the study and interpretation of the data. All authors have read and approved the manuscript.

