## Supplemental figures for "GPNMB overexpression- a marker of resistance to CDK4/6 inhibitors"

#### Supplementary Fig. S1

**A**

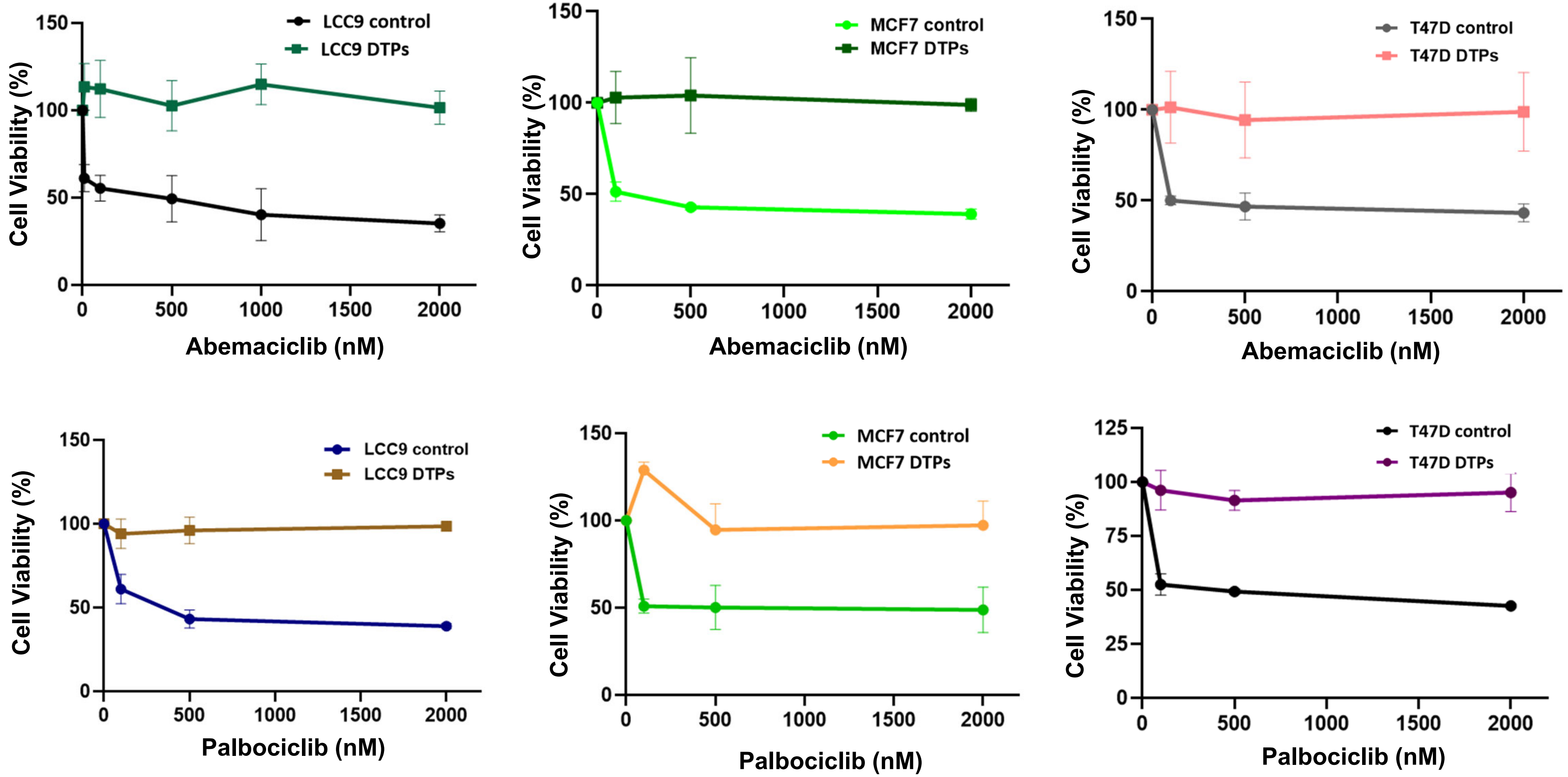

### Supplementary Fig. S1

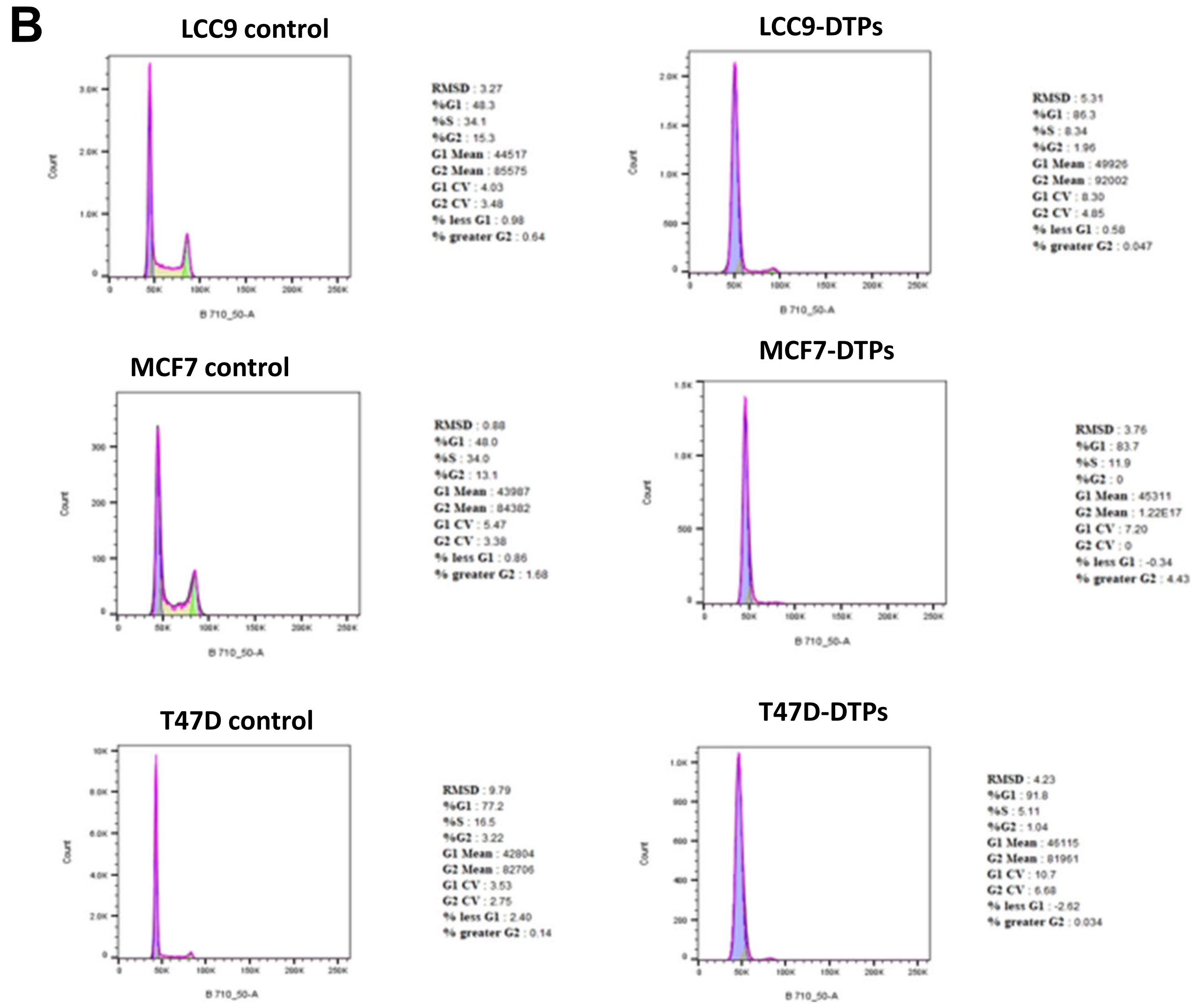


#### Abemaciclib

| | Cell Cycle G1% | Cell Cycle S% | Cell Cycle G2% |
|---|---|---|---|
| LCC9 | 48.3 | 34.1 | 15.3 |
| LCC9 DTPs | 86.3 | 8.34 | 1.96 |
| MCF7 | 48 | 34 | 13.1 |
| MCF7 DTPs | 83.7 | 11.9 | 0.2 |
| T47D | 77.2 | 16.5 | 3.22 |
| T47D DTPs | 91.8 | 5.11 | 1.04 |

#### Supplementary Fig. S1

##### LCC9-control

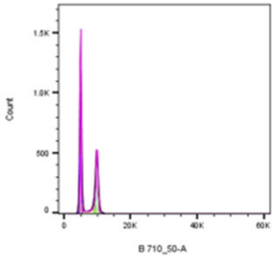


RMSD : 2.38
%G1 : 48.2
%S : 14.6
%G2 : 30.8
G1 Mean : 4950
G2 Mean : 9817
G1 CV : 3.37
G2 CV : 3.66
% less G1 : 2.46
% greater G2 : 2.24

##### LCC9 DTPs

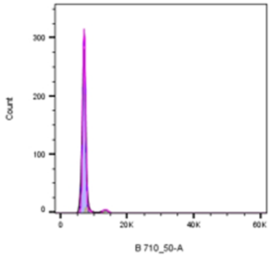


RMSD : 0.90
%G1 : 90.1
%S : 6.47
%G2 : 2.66
G1 Mean : 6993
G2 Mean : 13535
G1 CV : 6.85
G2 CV : 6.43
% less G1 : 2.81
% greater G2 : 9.80E-3

##### MCF7-control

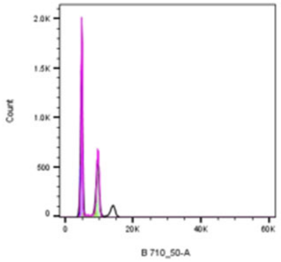


RMSD : 6.54
%G1 : 53.3
%S : 6.57
%G2 : 28.7
G1 Mean : 4800
G2 Mean : 9541
G1 CV : 4.59
G2 CV : 4.28
% less G1 : 2.13
% greater G2 : 11.4

##### MCF7 DTPs

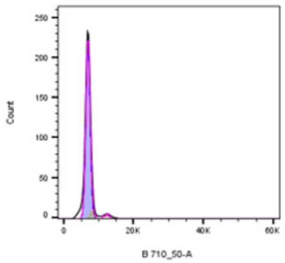


RMSD : 0.62
%G1 : 85.9
%S : 8.54
%G2 : 0
G1 Mean : 6854
G2 Mean : 2.50E8
G1 CV : 10.3
G2 CV : 0
% less G1 : 4.35
% greater G2 : 0

##### T47D-control

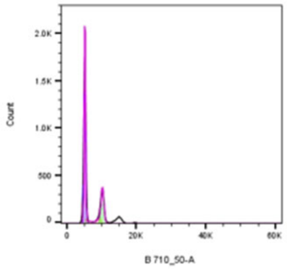


RMSD : 6.22
%G1 : 62.6
%S : 7.92
%G2 : 20.4
G1 Mean : 5099
G2 Mean : 10160
G1 CV : 4.54
G2 CV : 4.66
% less G1 : 2.42
% greater G2 : 9.92

##### T47D DTPs

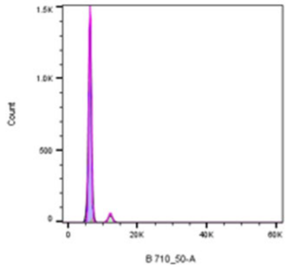


RMSD : 2.40
%G1 : 90.8
%S : 2.20
%G2 : 5.50
G1 Mean : 6261
G2 Mean : 12109
G1 CV : 6.42
G2 CV : 5.18
% less G1 : 2.65
% greater G2 : 0.37

##### Palbociclib

| | Cell Cycle G1% | Cell Cycle S% | Cell Cycle G2% |
|---|---|---|---|
| LCC9 | 48.2 | 14.6 | 30.8 |
| LCC9 DTPs | 90.1 | 6.47 | 2.66 |
| MCF7 | 53.3 | 6.57 | 28.7 |
| MCF7 DTPs | 85.9 | 8.54 | 0 |
| T47D | 62.6 | 7.92 | 20.4 |
| T47D DTPs | 90.8 | 2.2 | 5.5 |

#### Supplementary Fig. S1

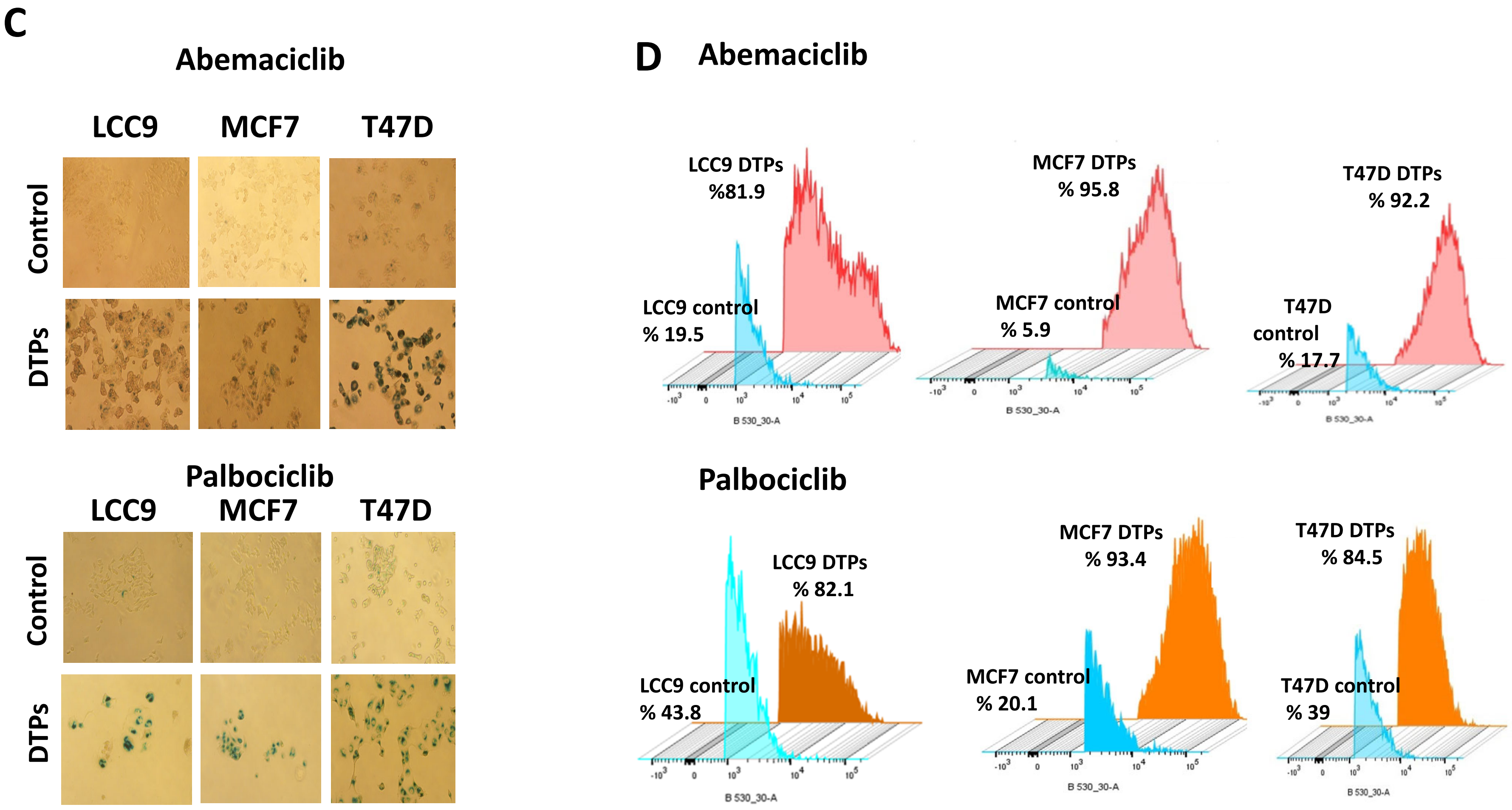

#### Supplementary Fig. S2

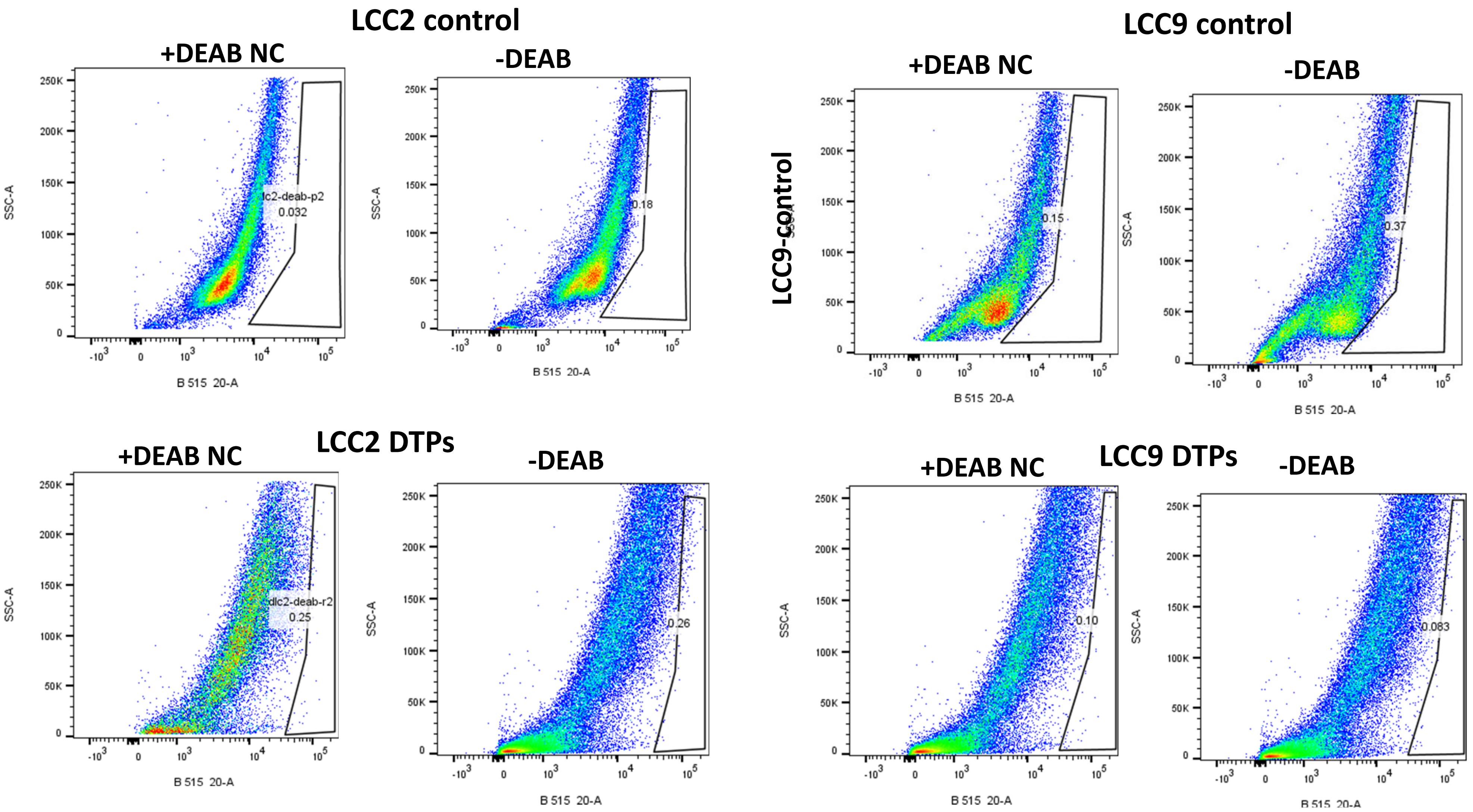

#### Supplementary Fig. S2

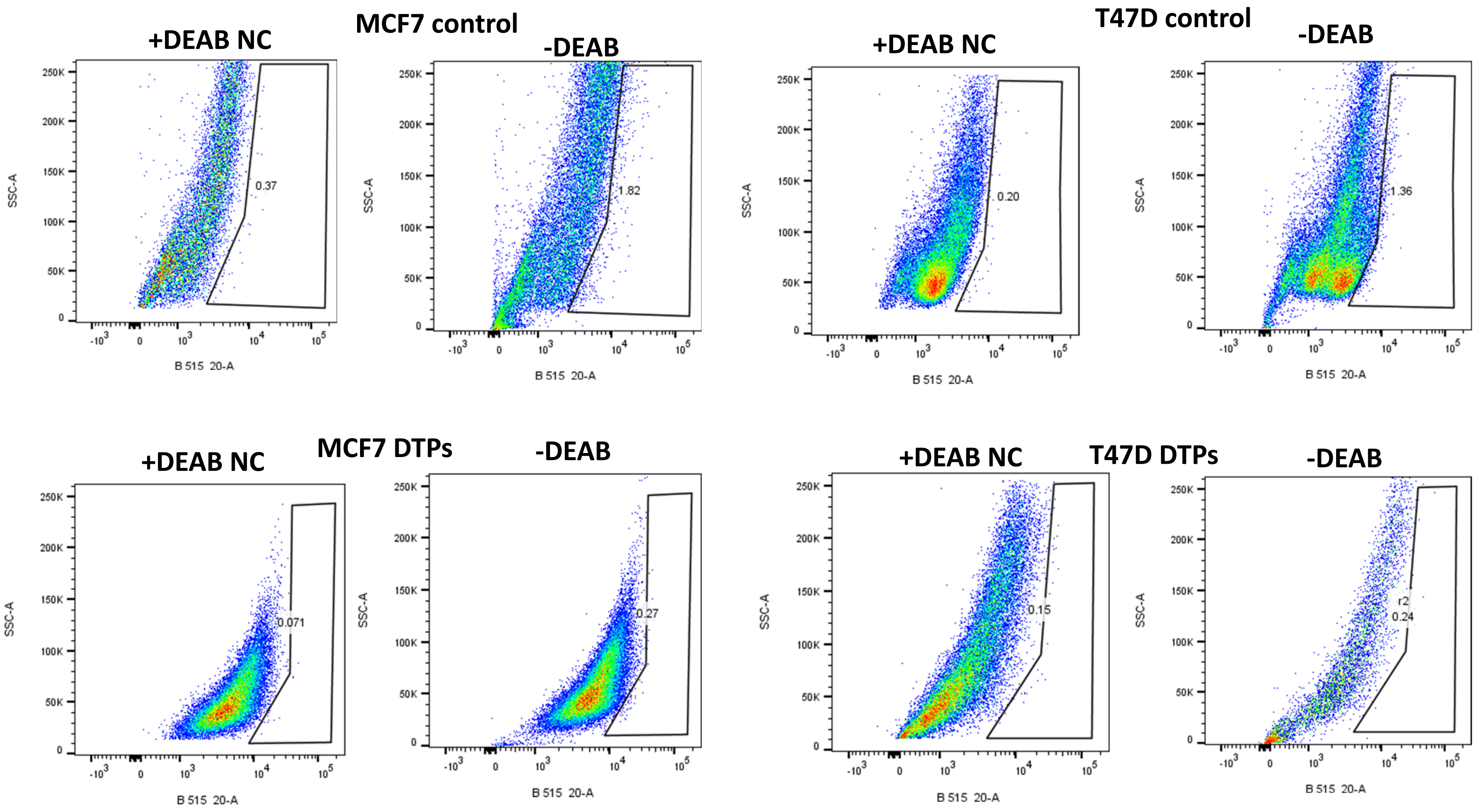

#### Supplementary Fig. S3

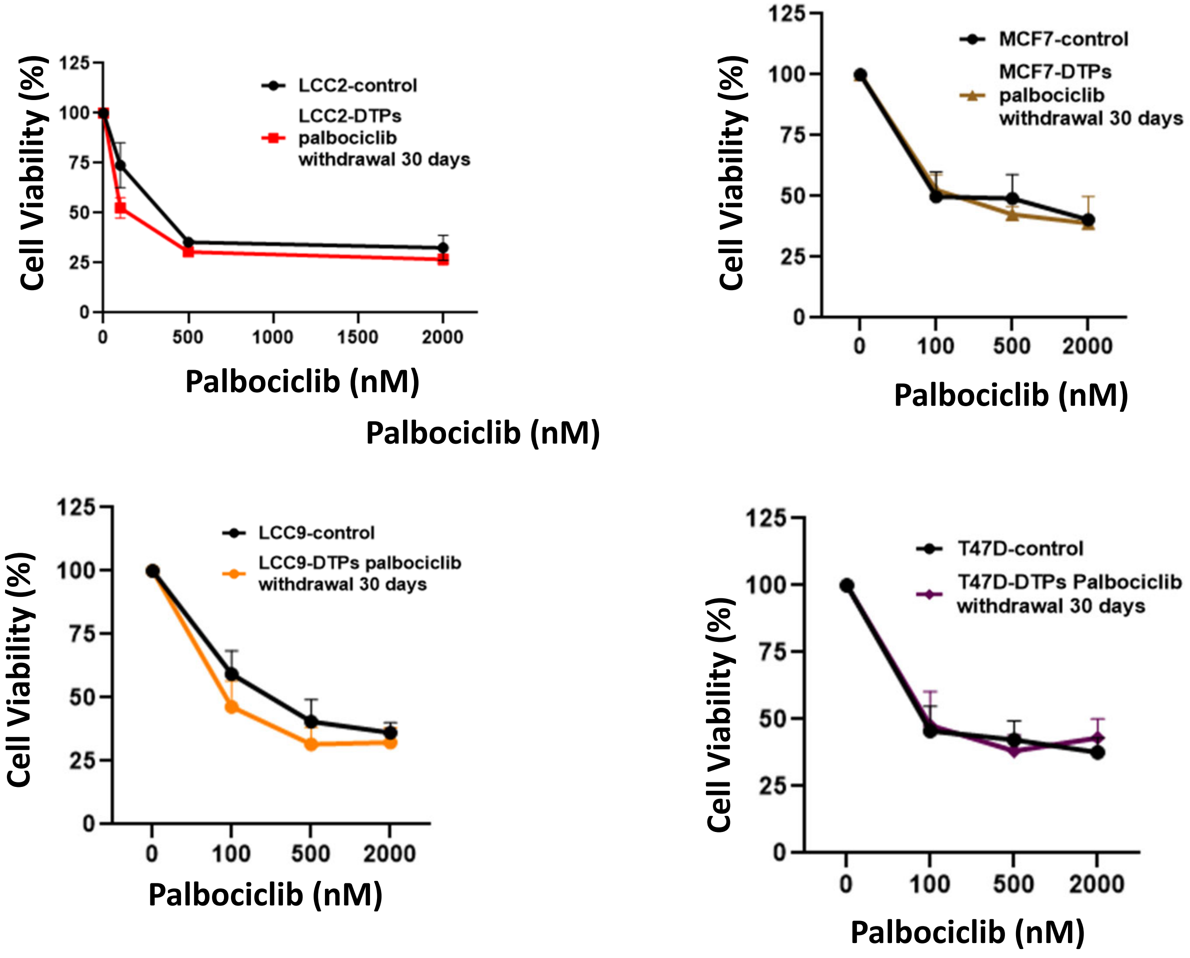

#### Supplementary Fig. S4

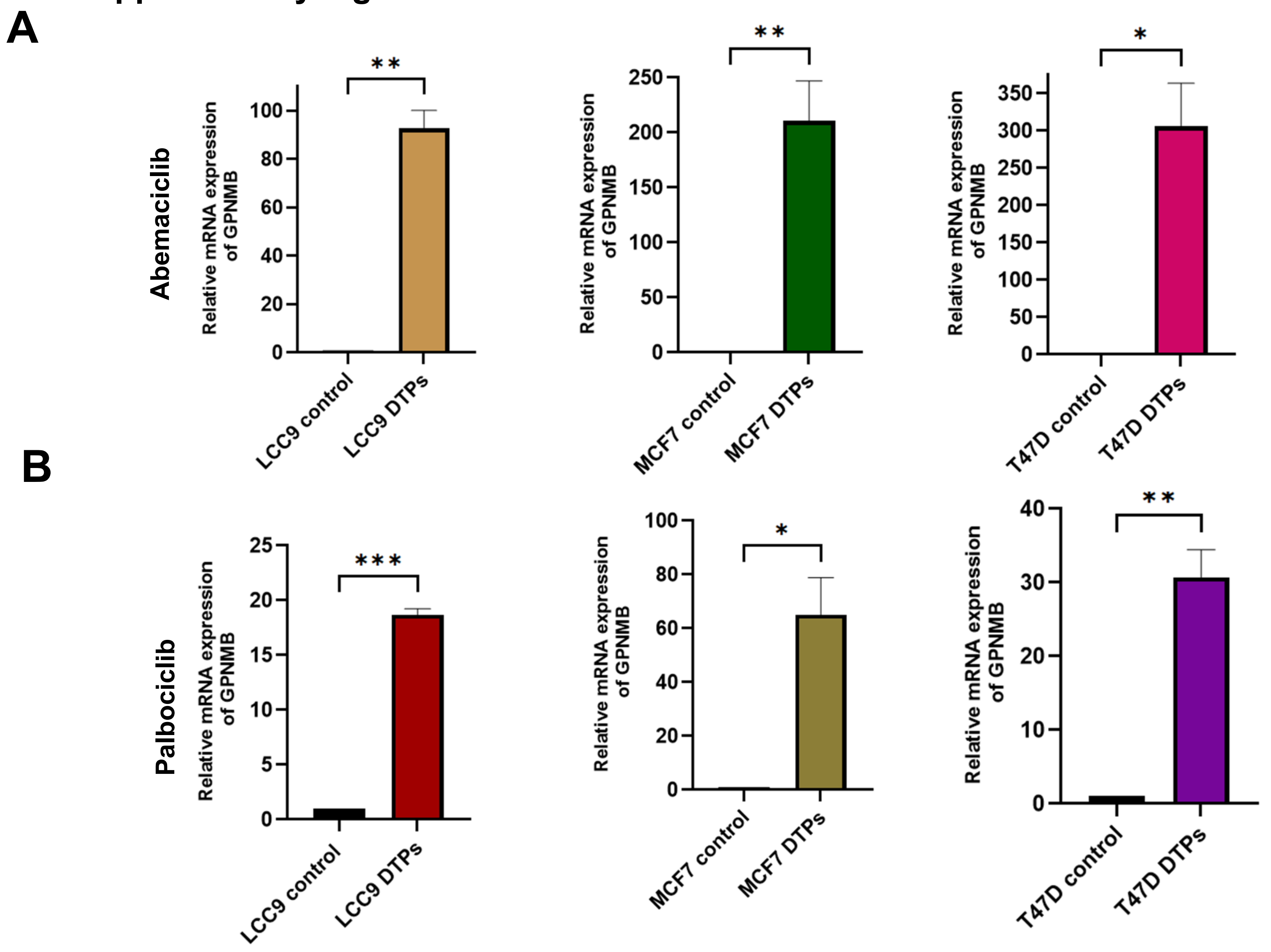

Supplementary Fig. S5

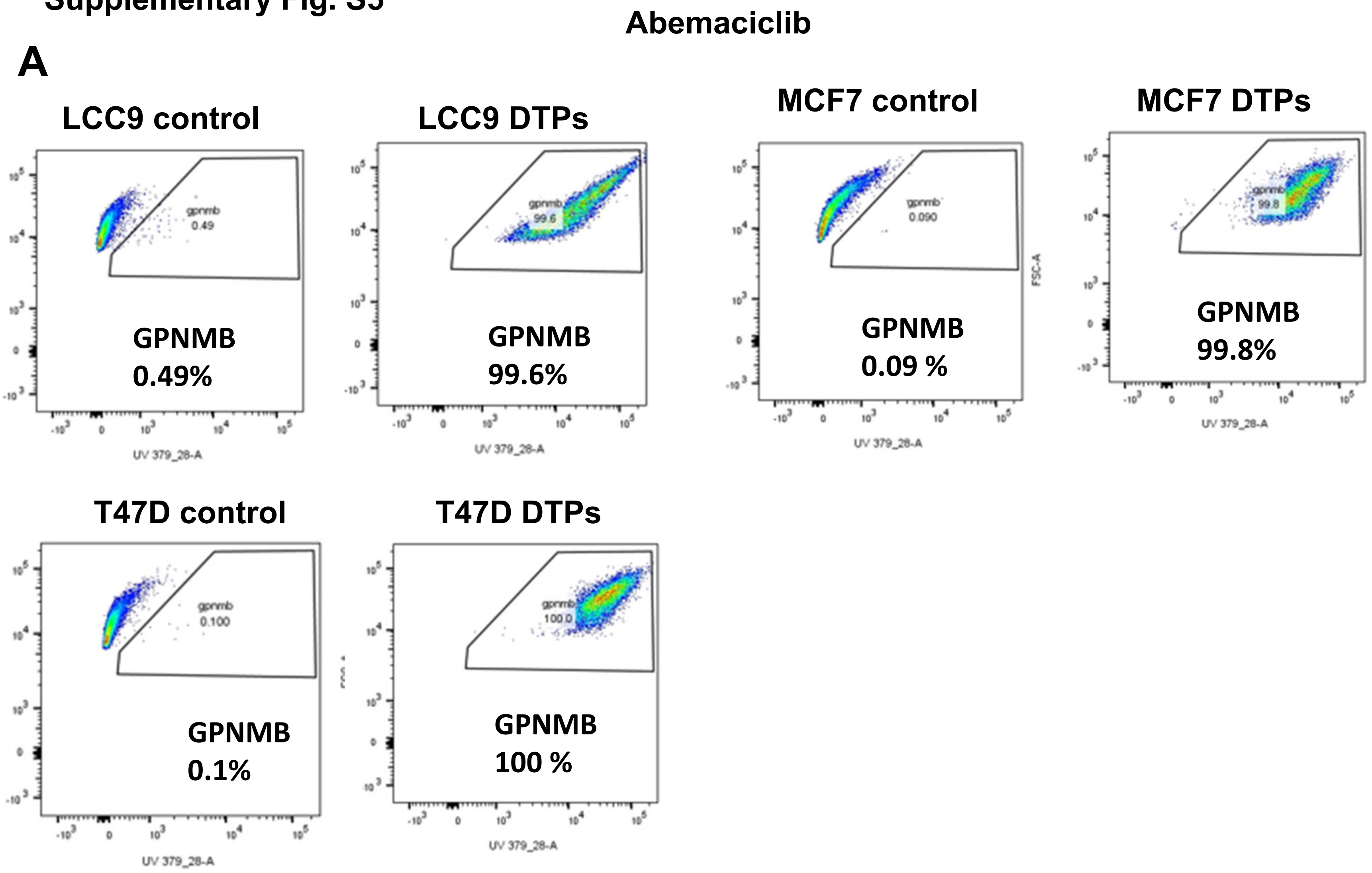

B

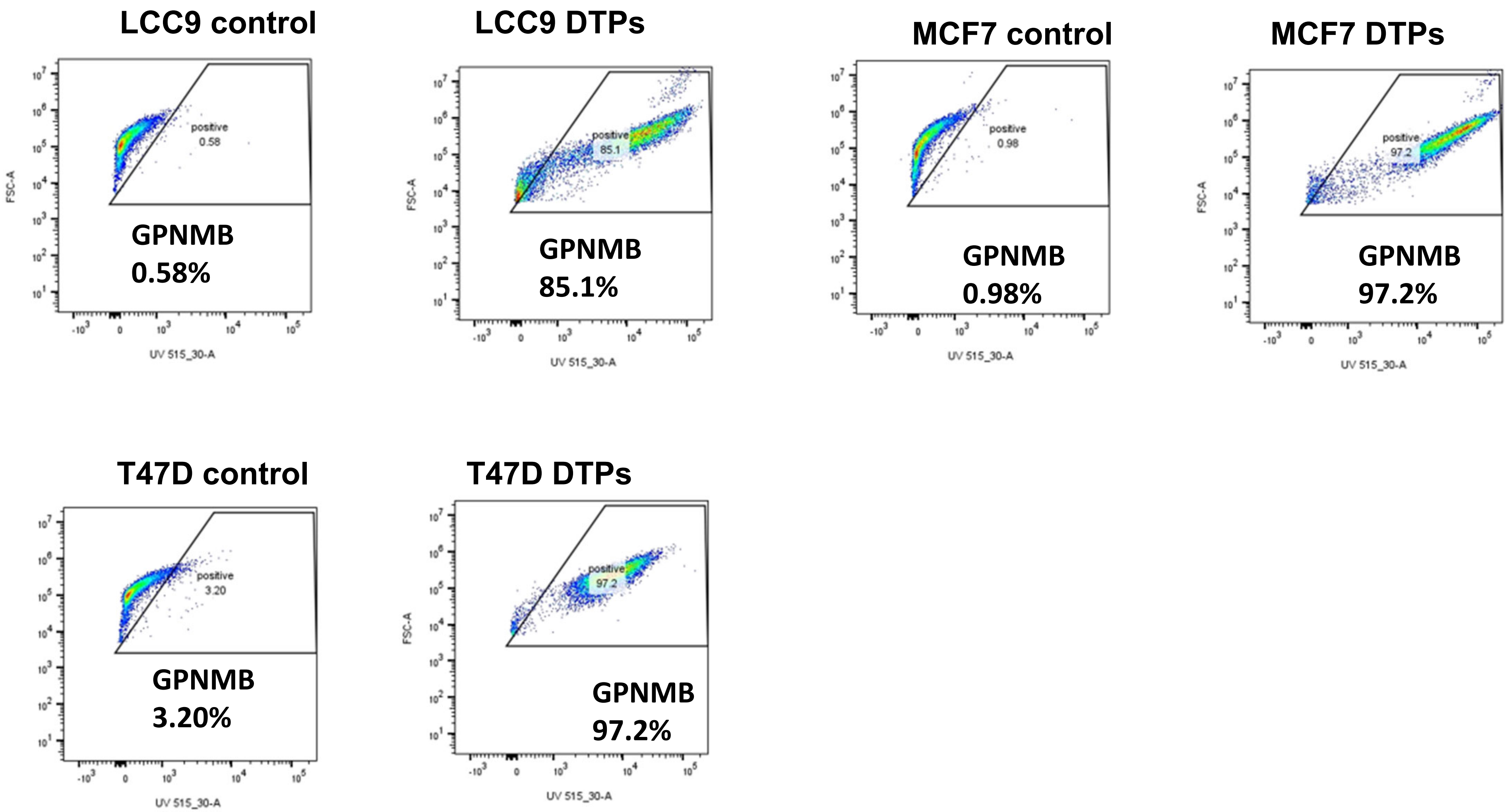
LCC9 control
positive
0.58
GPNMB
0.58%
LCC9 DTPs
positive
85.1
GPNMB
85.1%
MCF7 control
positive
0.98
GPNMB
0.98%
MCF7 DTPs
positive
97.2
GPNMB
97.2%
T47D control
positive
3.20
GPNMB
3.20%
T47D DTPs
positive
97.2
GPNMB
97.2%
FSC-A
UV 515_30-A

**Supplementary Fig. S6**

**spheroid immunofluorescent assay**

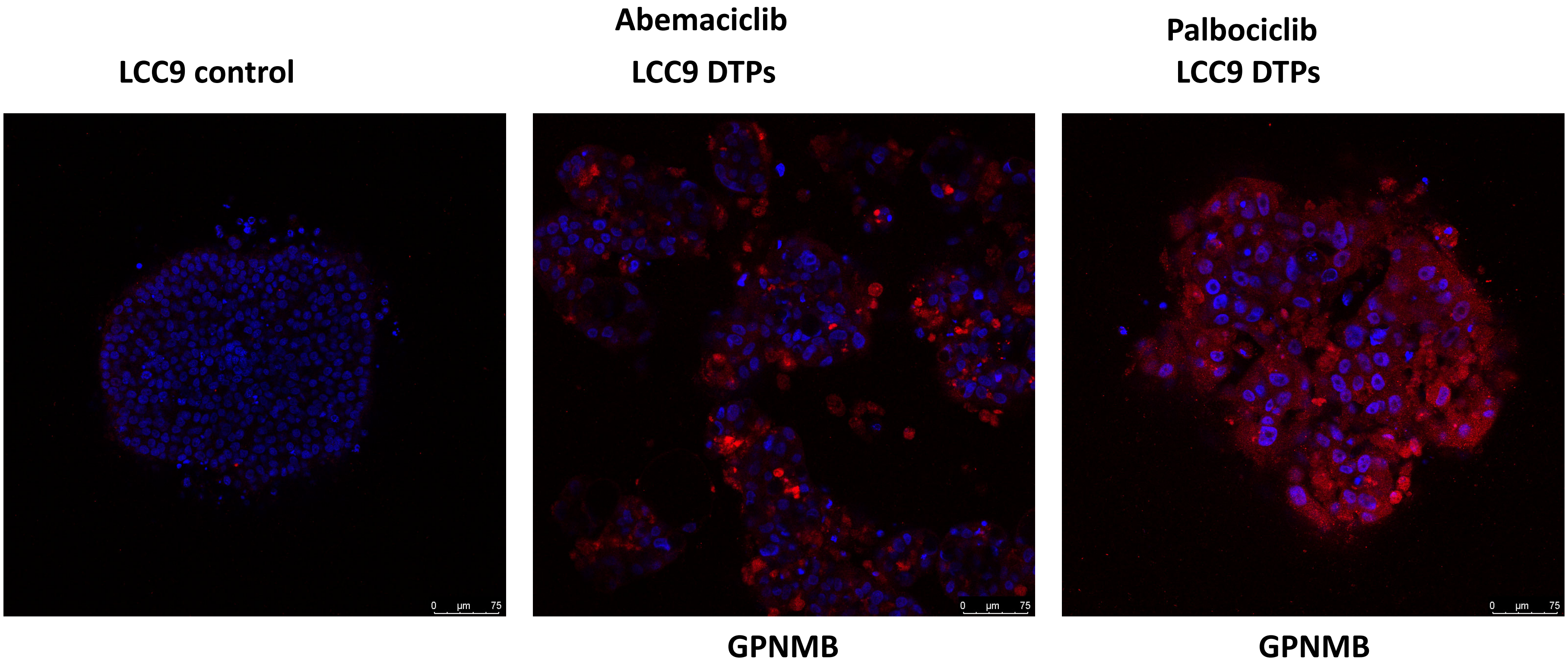

**Supplementary Fig. S6**

**spheroid immunofluorescent assay**

**MCF7 control** | **Abemaciclib MCF7 DTPs** | **Palbociclib MCF7 DTPs**

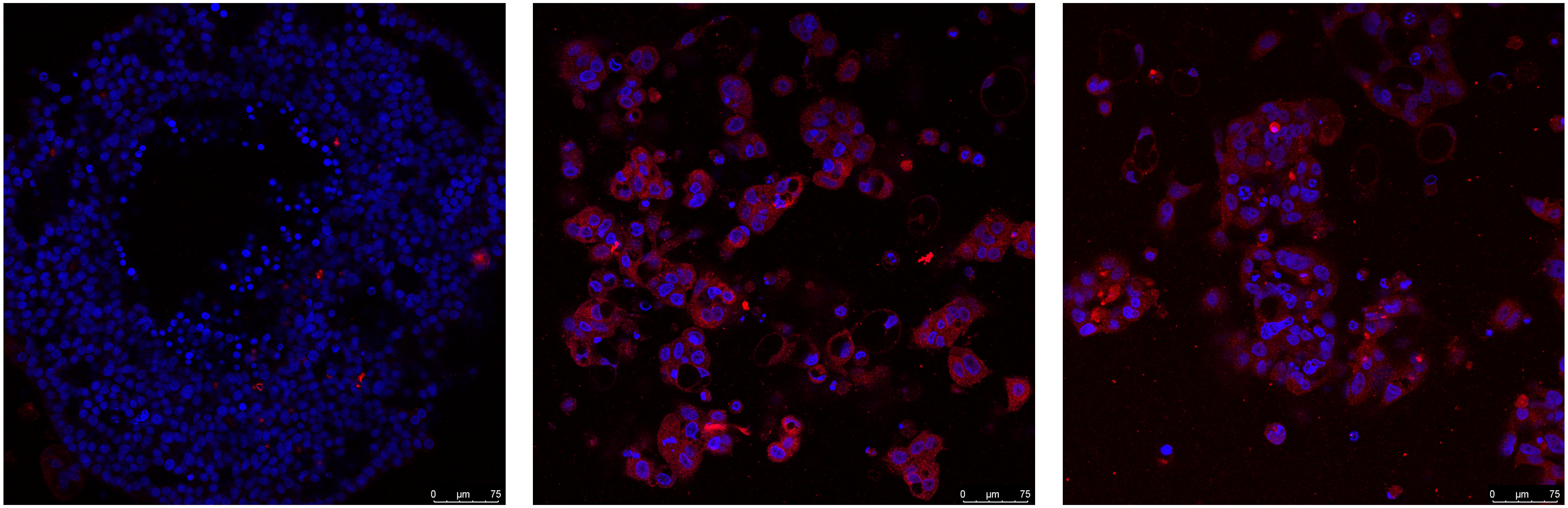


**GPNMB** | **GPNMB**

#### Supplementary Fig. S6

**spheroid immunofluorescent assay**

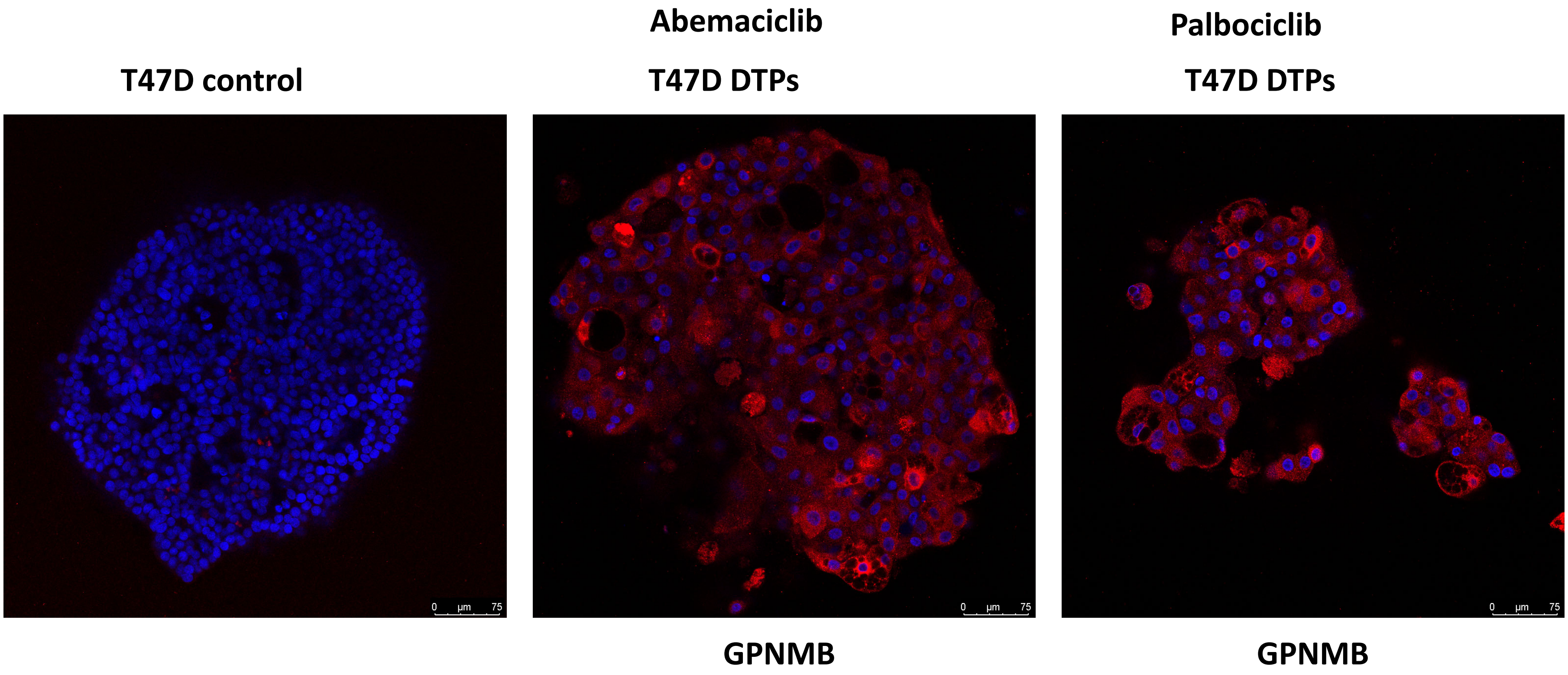

#### Supplementary Fig. S7

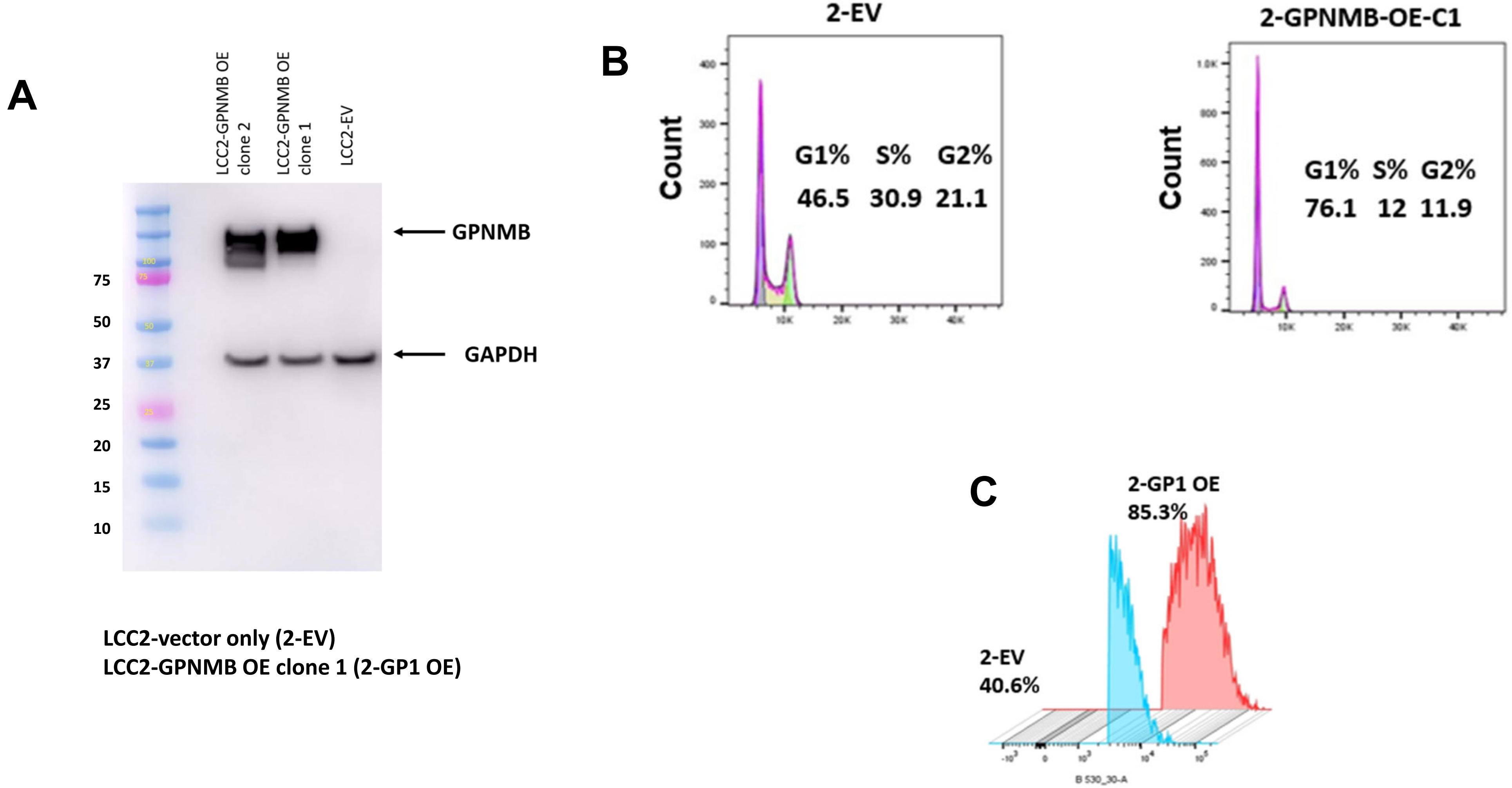


LCC2-vector only (2-EV)
LCC2-GPNMB OE clone 1 (2-GP1 OE)

**Supplementary Fig. S8**

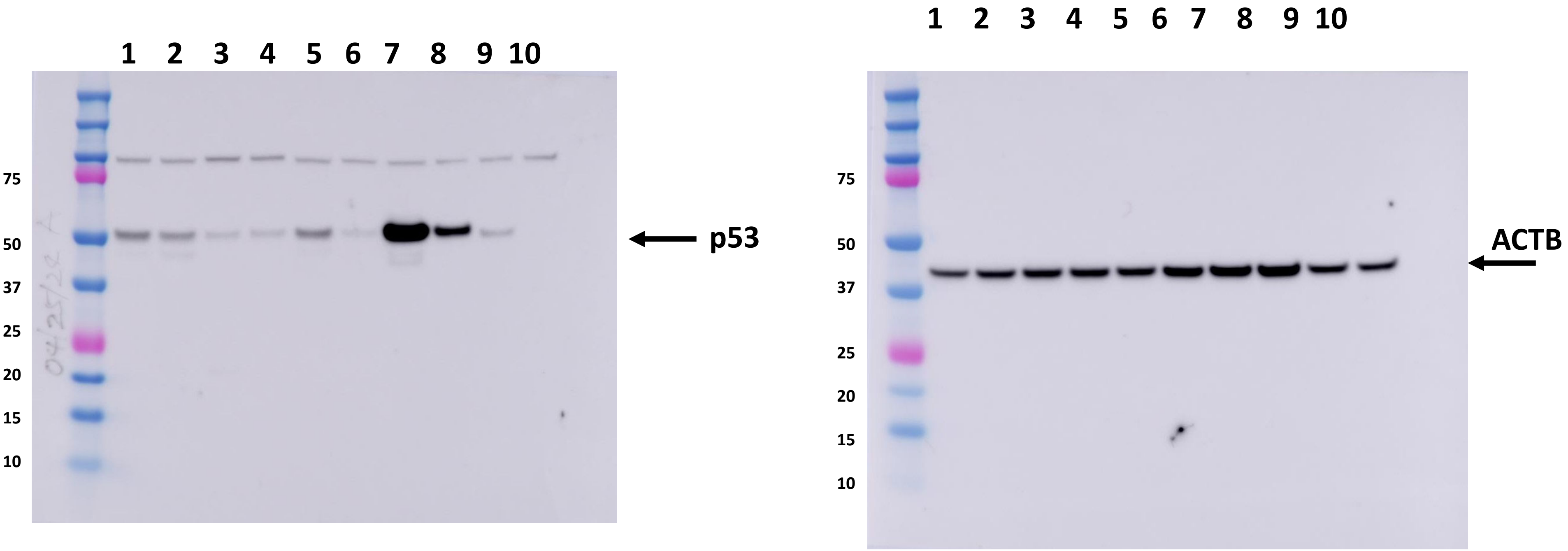


Samples**: # 1** LCC2 control**, #2** LCC2-DTPs**; #3** LCC9 control**, #4** LCC9-DTPs; **#5** MCF-7 control, **#6** MCF-7-DTPs; **#7** T47D control, **#8** T47D-DTPs; **#9** ZR75 control, **#10** ZR75-DTPs
